# Phosphoproteomics of Arabidopsis Highly ABA-Induced1 identifies AT-Hook Like10 phosphorylation required for stress growth regulation

**DOI:** 10.1101/413013

**Authors:** Min May Wong, Govinal Badiger Bhaskara, Tuan-Nan Wen, Wen-Dar Lin, Thao Thi Nguyen, Geeng Loo Chong, Paul E. Verslues

## Abstract

The Clade A protein phosphatase 2C Highly ABA-Induced 1 (HAI1) plays an important role in stress signaling yet little information is available on HAI1-regulated phosphoproteins. Quantitative phosphoproteomics identified phosphopeptides of increased abundance in *hai1-2* in unstressed plants and in plants exposed to low water potential (drought) stress. The identity and localization of the phosphoproteins as well as enrichment of specific phosphorylation motifs indicated that these phosphorylation sites may be regulated directly by HAI1 or by HAI1-regulated kinases including Mitogen-Activated Protein Kinases (MPKs), Sucrose-non fermenting Related Kinase 2 (SnRK2s) or Casein Kinases. One of the phosphosites putatively regulated by HAI1 was S313/S314 of AT Hook-Like10 (AHL10), a DNA binding protein of unclear function. HAI1 could directly dephosphorylate AHL10 *in vitro* and the level of *HAI1* expression affected the abundance of phosphorylated AHL10 *in vivo.* AHL10 S314 phosphorylation was critical for restriction of plant growth under low water potential stress and for regulation of Jasmonic Acid and Auxin-related gene expression as well as expression of developmental regulators including *Shootmeristemless* (*STM*). These genes were also mis-regulated in *hai1-2*. AHL10 S314 phosphorylation was required for AHL10 complexes to form foci within the nucleoplasm, suggesting that S314 phosphorylation may control AHL10 association with the nuclear matrix or with other transcriptional regulators. These data identify a set of HAI1-affected phosphorylation sites, show that HAI1-regulated phosphorylation of AHL10 S314 controls AHL10 function and localization and also indicate that HAI1-AHL10 signaling coordinates growth with stress and defense responses.

## Introduction

Many types of environmental stress, both abiotic and biotic, limit plant growth. Drought stress is of particular interest and challenge because of its effect on crop productivity as well as our lack of understanding of the mechanisms plants use to sense and respond to soil drying and reduced water potential (ψ_w_) during drought. Low ψ_w_.-induced accumulation of Abscisic Acid (ABA) controls downstream responses including growth and gene expression (1, 2). Plants perceive increased ABA levels via a pathway consisting of the PYR/PYL/RCAR ABA receptors (hereafter referred to as PYLs), Clade A protein phosphatase 2Cs (PP2Cs) and Sucrose Non-Fermenting Related Kinase 2 (SnRK2) protein kinases (3, 4). As ABA increases, formation of PYL-ABA-PP2C complexes inhibits PP2C activity. This releases suppression of SnRK2 activity by allowing the SnRK2s to autophosphorylate and phosphorylate downstream target proteins. Proteins regulated by SnRK2-mediated phosphorylation include nuclear and plasma membrane localized proteins. PYL-PP2C signaling can regulate Mitogen-activated Protein Kinase (MPK) signaling cascades which are active in ABA and abiotic stress signaling (5) in addition to their roles in defense signaling. Identification of phosphorylation sites regulated by SnRK2s, MPKs and ABA signaling is an area of active research (6–10). Despite the success of these studies in identifying kinase targets and stress-affected phosphorylation sites, phosphoproteomics has yet to be applied to identify regulatory targets of the Clade A PP2Cs. We are aware of only one phosphoproteomic study of plant PP2Cs, an analysis of Clade E PP2Cs (11).

Six of the nine Clade A PP2Cs were identified in forward genetic screens for altered ABA sensitivity of seed germination (see for example 12–14). Interestingly, the remaining three Clade A PP2Cs, Highly ABA-Induced1 (HAI1), AKT-Interacting Protein1 (AIP1)/HAI2, and HAI3, were not identified in such screens as their effect on ABA sensitivity of seed germination is less (15, 16). Gene expression of the *HAI* PP2Cs is strongly induced by ABA as well as drought and salt stress (15, 17). *HAI* PP2C mutants, and *hai1* in particular, had a strong effect on osmotic adjustment and maintenance of fresh weight at low ψ_w_ (15). The three HAI PP2Cs are not regulated by Enhancer of ABA co-Receptor 1 (EAR1), which promotes the activity of the six other Clade A PP2Cs and thereby affects ABA sensitivity (18). Also, Mine et al. (19) found that the HAI PP2Cs, but not the Clade A PP2C ABI2, could interact with MPK3 and MPK6. They also demonstrated that HAI1 could directly dephosphorylate these MPKs. *HAI1* was induced by coronatine-mediated activation of the JA-signaling factor MYC2 and HAI1 suppression of MPK3 and MPK6 activation promoted virulence of *Pseudomonas syringae* (19). *HAI1* was also identified as a target of the transcription factor SCARECROW, a root development regulator (20). Together these studies indicate that there are both overlapping aspects and diversification among the Clade A PP2Cs in terms how strongly they affect specific stress and ABA-related phenotypes and how their phosphatase activity is regulated. The data also suggest a prominent role for HAI1 in coordination of abiotic stress, development, and defense responses

We used quantitative phosphoproteomics of *hai1-2* to identify phosphorylation sites putatively regulated by HAI1. One of the proteins with increased phosphopeptide abundance in *hai1-2* was AT-Hook Like10 (AHL10). Phosphorylation of AHL10 at the site affected by HAI1 was required for AHL10 to suppress growth, regulate expression of hormone and development related genes at low ψ_w_ and localize to foci within the nucleoplasm. These data indicate that HAI1 and AHL10 connect stress signaling to growth and developmental regulation.

## Results

### HAI1-affected phosphorylation sites identified by quantitative phosphoproteomics

Longer term low ψ_w_ stress strongly induces *HAI1* expression (15). Thus, quantitative phosphoproteomics of wild type and *hai1-2* was performed for seedlings maintained at high ψ_w_ (unstressed control) and for seedlings transferred to low ψ_w_ (−1.2 MPa) for 96 h. The analysis of *hai1-2* (Dataset S1) was conducted in the same set of iTRAQ labeling experiments as the wild type and Clade E Growth Regulating (EGR) PP2C data previously reported by our laboratory (11, 21). The criteria used to define phosphopeptides of altered abundance in *hai1-2* versus wild type in either control or stress treatment were fold change ≥ 1.5 and P ≤ 0.05. Calculation of q values to estimate false discovery rates showed that P = 0.05 corresponded to q = 0.11 for the control data and q = 0.14 for the stress data. In the unstressed control these criteria identified 61 phosphopeptides from 56 proteins that were significantly more abundant in *hai1-2* compared to wild type while only 18 phosphopeptides from 17 proteins were significantly less abundant in *hai1-2* (Dataset S2). Similarly, at low ψ_w_ 40 phosphopeptides from 39 proteins were significantly more abundant in *hai1-2* while 8 phosphopeptides from 7 proteins were less abundant (Dataset S3). Even proteins with highly increased phosphopeptide abundance had no significant change in gene expression in *hai1-2* (Fig. 1A, the *hai1-2* transcriptome data has been discussed previously (15)). Thus, phosphoproteomics identified a different set of HAI1-affected loci than transcriptome analysis. Four proteins had increased phosphopeptide abundance in both control and stress treatments (SI Appendix Fig. S1). While this limited overlap could indicate that HAI1 regulates different sets of proteins in the control and stress conditions, the incomplete coverage of the phosphoproteome should also be kept in mind.

**Figure 1:**
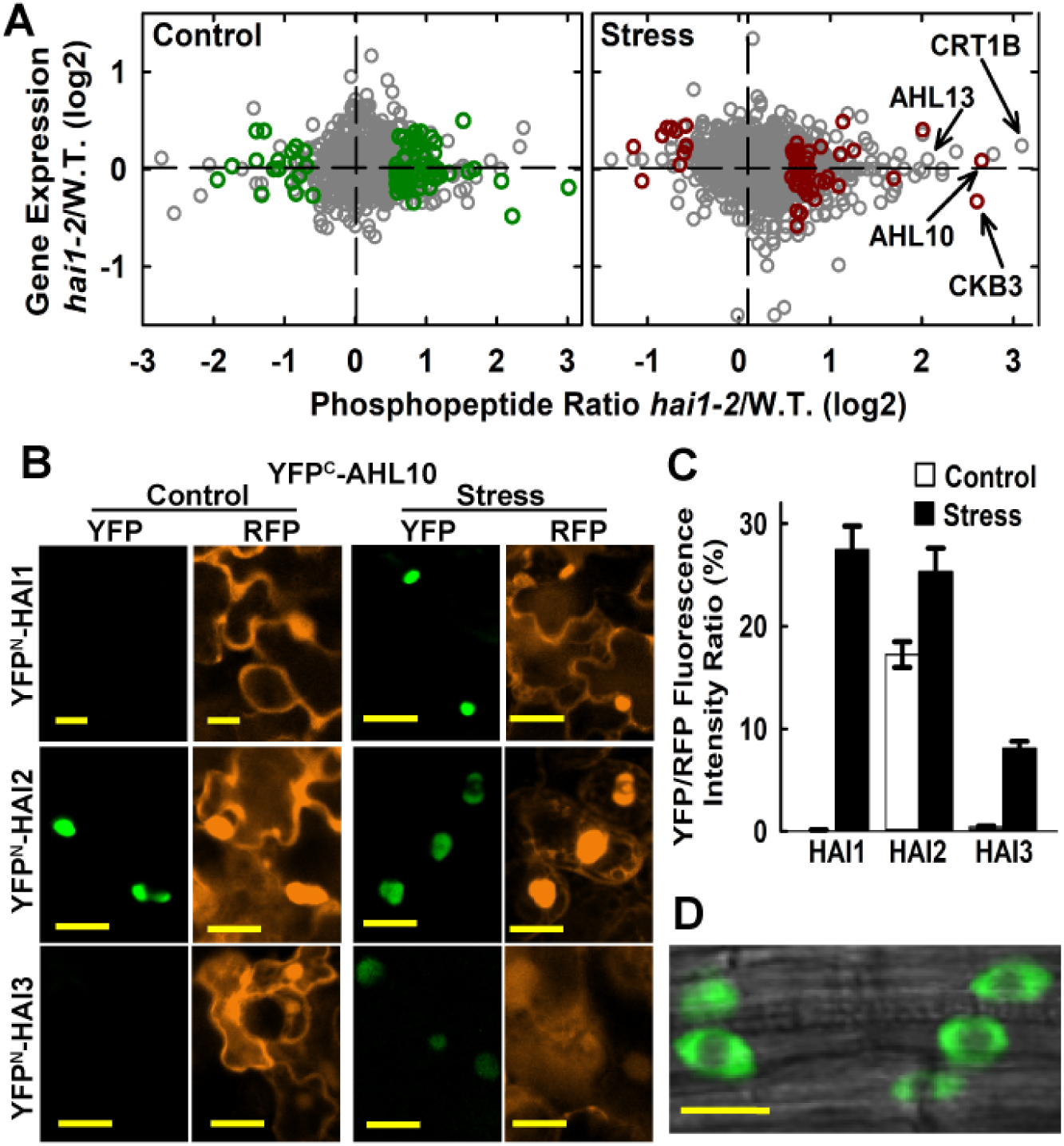
Phosphoproteomics of *hai1-2* identifies a set of HAI1-affected phosphoproteins, including AHL10. A. Phosphopeptide abundance versus gene expression for *hai1-2* compared to wild type. Dark green or red symbols indicate phosphopeptides with significantly increased or decreased abundance (P ≤0.05 by one sample T-test and fold change ≥ 1.5) in *hai1-2* compared to wild type for control and stress (−1.2 MPa, 96 h) treatments (Dataset S2, S3). Other phosphopeptide data are plotted using gray symbols (Dataset S1). Transcriptome analysis of *hai1-2* has been previously described (15). B. Ratiometric Bimolecular Fluorescence Complementation (rBiFC) assays of HAI PP2C interaction with AHL10. Interactions were tested by transient expression in Arabidopsis seedlings under unstressed control conditions or after transfer to −1.2 MPa for 48 h before imaging. RFP panels show fluorescence of the constitutively expressed RFP reporter was used to normalize the YFP fluorescence. Scale bars indicate 20 μm. C. Relative quantification of rBiFC interactions shown in B. The mean fluorescence intensity was measured for individual cells and the ratio of YFP to RFP intensity calculated. Data are ± SE (n = 20-25) combined from two independent experiments. D. AHL10 localization in plants expressing *AHL10*_*promoter*_:*AHL10-YFP* in the *ahl10-1* mutant background. Cells in root tip of unstressed seedlings are shown. An essentially identical localization pattern was observed in stress treated seedlings. Scale bar indicates 10 μm.

The *hai1-2* affected phosphorylation sites were enriched in several specific phosphorylation motifs (SI Appendix Fig. S2; Dataset S4). These included the [pSP] motif, consistent with phosphorylation by MPKs and other proline-directed kinases (10) as well as variations of the [RxxpS] motif which can be targeted by SnRKs and Calcium Dependent Protein Kinases (CPKs) (22) and serine surrounded by acidic residues (E or D) which may be recognized by Casein kinase II (23). Consistent with these enriched motifs, five of the putative HAI1 target proteins are also putative SnRK2 substrates (Dataset S5) (6, 7, 24). Also consistent with enrichment of the [RxxpS] motif, a phosphopeptide from CPK9 was more abundant in *hai1-2* (Dataset S2), suggesting that HAI1 may affect some [RxxpS] sites via regulation of CPKs as well as SnRK2s. Similarly, the proteins with phosphopeptides increased in *hai1-2* included three known MPK substrates, PHOS32 (25), SCL30 (26) and AT1G80180 (10), as well as 12 putative MPK substrates (Dataset S5) (9, 27). In relation to Casein kinases, a phosphopeptide from Casein Kinase 2 Beta Chain3 (CKB3) was increased in *hai1-2* under stress (Figure 1A) and a phosphopeptide from Casein Kinase1-Like Protein2 (CKL2) was increased in *hai1-2* in the control (Dataset S2). This was consistent with the enrichment of casein kinase phosphorylation motifs and observations that CKL2 affects ABA response (28, 29).

The proteins with increased phosphopeptide abundance in *hai1-2* under stress had a high prevalence of nuclear and plasma membrane localized proteins (SI Appendix Fig. S3). This was consistent with predominant localization of HAI1 in the nucleus but also partially along the plasma membrane (15, 30). Several of the proteins with increased phosphopeptide abundance in *hai1-2* have been previously found to be ABA or stress regulated. These include AKS1, AtSIK1, Annexin1 and the phosphatidic acid binding protein PLDRP1 (31). Interestingly several mRNA splicing-related proteins (RSP31, SCL30, RSZp22, RSZp33) were affected by *hai1-2* (Dataset S2, S3), consistent with previous reports that splicing protein phosphorylation is altered by stress or ABA and may be regulated by SnRK2 kinases (6, 7). It should be noted that our 96 h stress treatment (96 h), was advantageous to allow induction of *HAI1* expression. However, because of this longer stress treatment changes in phosphopeptide abundance could be influenced by changes in protein abundance as well as change in phosphorylation stoichiometry. While keeping this caveat in mind, the *hai1-2* phosphoproteomics dataset is overall a useful resource to uncover signaling mechanisms influenced by HAI1.

### AT-Hook Like10 (AHL10) interacts with and is dephosphorylated by HAI1

Bimolecular flourescence complementation (BiFC) assays were used as an initial screen for proteins which could be directly targeted by HAI1 (SI Appendix Fig. S4A). HAI1 interacted with AHL10 and CKB3, which had large increases in phosphopeptide abundance in *hai1-2* at low ψ_w_ (Fig. 1A), as well as a bZIP transcription factor (AT2G31370) which had a putative, but not significant, increase in phosphopeptide abundance (Dataset S1). Conversely, we did not see interaction with Calreticulin 1B (CRT1B, AT1G09210), which had the largest fold increase of all phosphopeptides but was variable and not significant (Fig. 1A), or PHOS32 and NF-YC11 (SI Appendix Fig. S4A).

We focused further attention on AHL10. Phosphopeptides with either AHL10 S313 or S314 phosphorylation were identified (SI Appendix Fig. S5). The AHL10 S313 phosphopeptide was strongly increased in *hai1-2* under stress (Fig. 1A, Dataset S3) while *AHL10* gene expression was only slightly increased by stress in wild type (SI Appendix Fig. S4B) and not affected by *hai1-2* (Fig. 1A). Note that there was some ambiguity in assigning the phosphorylation site to S313 or S314 in both of the AHL10 phosphopeptides identified. The same peptide with putative S313/S314 phosphorylation was identified in multiple previous studies (6, 7, 26, 32–38). Interestingly, a phosphopeptide from AHL13 (AT4G17950) was also putatively (but not significantly) increased in *hai1-2* at low ψ_w_ (Fig. 1A, Dataset S1, SI Appendix Fig. S6). AHL10 and AHL13 are closely related Clade B AHLs (SI Appendix Fig.S7A) and the AHL10 S314 and AHL13 phosphorylation sites are in equivalent positions in the C-terminal region of both proteins (SI Appendix Fig. S7B). Other Clade B AHLs lack this phosphorylation site.

Ratiometric BiFC (rBiFC) comparison of relative interaction intensity found that HAI1-AHL10 interaction was promoted by low ψ_w_ (Fig. 1B, C). AHL10 also interacted with HAI2 and HAI3. The HAI PP2C-AHL10 interaction occurred in a diffuse pattern in the nucleoplasm (Fig. 1B). A similar diffuse localization in the nucleoplasm and exclusion from nucleolus was seen in transgenic plants expressing *AHL10*_*promoter*_:*AHL10-YFP* in the *ahl10-1* background (Fig. 1D).

To test whether HAI1 could directly dephosphorylate AHL10, AHL10-YFP was immunoprecipitated from stress treated (−1.2 MPa) *AHL10*_*promoter*_:*AHL10-YFP/ahl10-1hai1-2* plants and analyzed on Phos-tag gels after *in vitro* phosphatase treatment (*AHL10* mutant and transgenic lines are described in SI Appendix Fig S8). In mock incubated samples (no phosphatase), multiple bands of phosphorylated AHL10 could be observed (indicated by asterisks in Fig. 2A), consistent with PhosPhAt listing of three experimentally observed AHL10 phosphorylation sites (S297, T311, S317) in addition to S313 and S314 (7, 33, 34, 36, 38). Incubation with non-specific phosphatase (Calf Intestinal Phosphatase, C.I.P.) completely dephosphorylated AHL10 (Fig. 2A). Treatment with recombinant HAI1 eliminated the most highly phosphorylated AHL10 bands (those having slowest migration on Phos-tag gel) but did not completely dephosphorylate AHL10 (Fig. 2A). This indicated that HAI1 could specifically dephosphorylate some, but not all, AHL10 phosphorylation sites.

**Figure 2:**
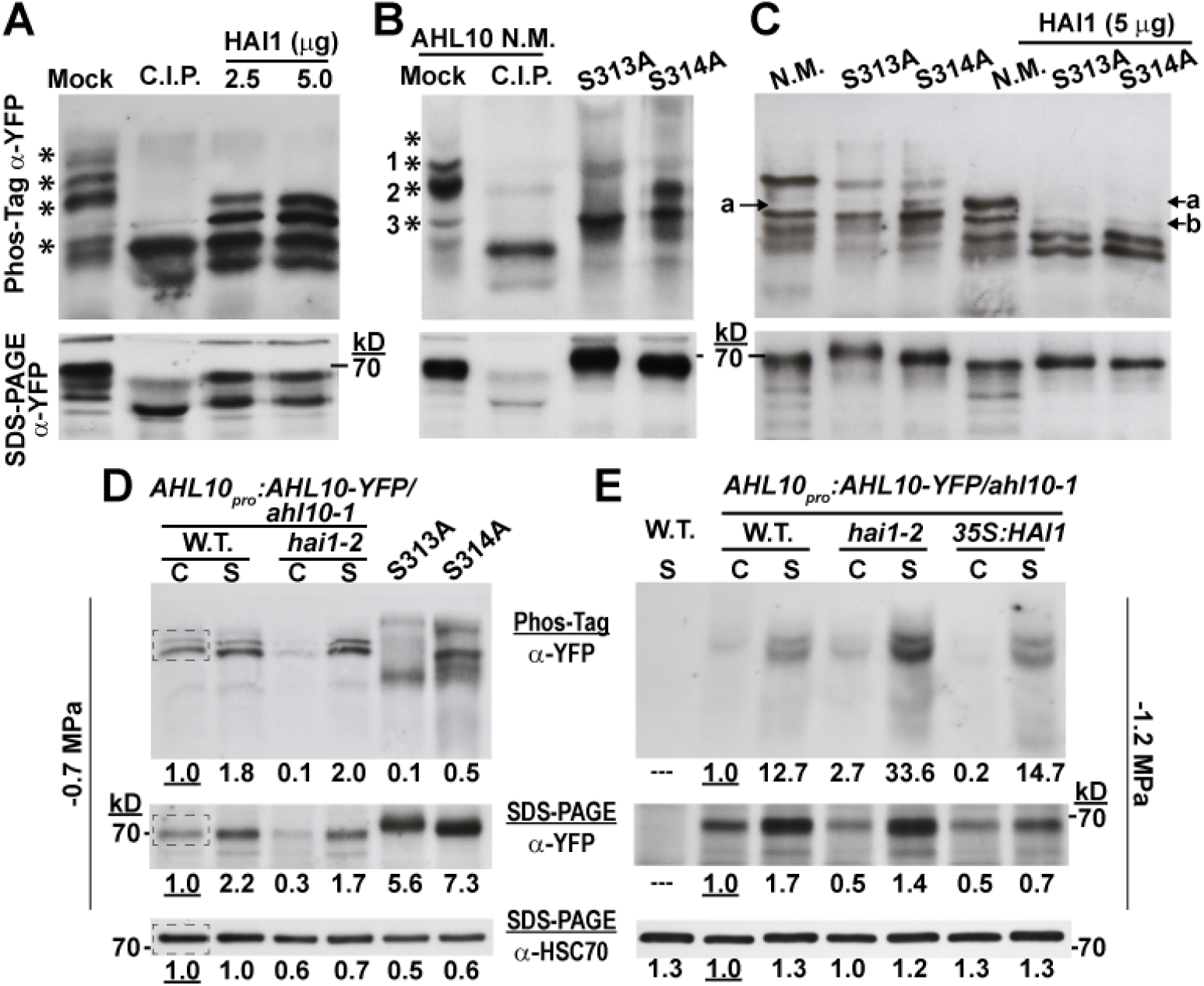
Phosphorylated AHL10 increases under stress and can be directly dephosphorylated by HAI1. A. AHL10-YFP immunoprecipitated from *AHL10*_*promoter*_:*AHL10-YFP/ahl10-1hai1-2* plants after exposure to −1.2 MPa stress was dephosphorylated using recombinant HAI1 or Calf Intestinal Phosphatase (C.I.P.). Aliquots of the same samples were run on Phos-tag gel or SDS-PAGE. Asterisks (*) along the Phos-tag blot indicate phosphorylated forms of AHL10 which were dephosphorylated (shifted down in the Phos-tag gel) by C.I.P. treatment. The expected molecular weight of AHL10-YFP is 69 kD. The experiment was repeated with consistent results. B. Phos-tag gel analysis of non-mutated AHL10 (N.M.) as well phosphonull AHL10 (AHL10^S313A^, AHL10^S314A^) immunoprecipitated from plants expressing *35S:YFP-AHL10* in the *ahl10-1* mutant background. Experimental conditions were the same as for A and asterisks mark bands of phosphorylated AHL10 present in the N.M. lane that are eliminated by C.I.P. treatment. The expected molecular weight of YFP-AHL10 is 70 kD. C. In vitro de-phosphorylation of non-mutated (N.M.) and phosphonull AHL10 (AHL10^S313A^, AHL10^S314A^) immunoprecipitated from plants expressing *35S:YFP-AHL10* in the *ahl10-1* mutant background. Experimental conditions were the same as in A except that less starting protein was used for the AHL10^S313A^ and AHL10^S314A^ immunoprecipitation compared to AHL10 N.M. (75 vs 100 μg, respectively) to ensure that similar amount of all AHL10 isoforms was used in the dephosphorylation assay. D. Phos-tag gel analysis of *AHL10*_*promoter*_::*AHL10-YFP/ahl10-1* and *AHL10*_*promoter*_::*AHL10-YFP/ahl10-1hai1-2* total protein extracted from seedlings in the control (C) and −0.7 MPa stress (S) treatments. Aliquots of the same samples were run on Phos-tag (top) and SDS-PAGE (middle) gels and band intensities of AHL10-YFP quantified. For comparison total protein extract from *35S:YFP-AHL10* S313A or S314A (in the *ahl10-1* mutant background) in the −0.7 MPa treatment was also analyzed. Each lane was loaded with 25 μg of protein. Blots from the SDS-PAGE separation were first probed with anti-YFP to detect AHL10 and then stripped and re-probed with anti-HSC70 as a loading control. The dashed box in the wild type control lanes indicates the region selected from each lane for quantification of band intensities. Band intensities relative to the wild type unstressed control are indicated by the numbers below each lane. Note that unphosphorylated AHL10 could not be resolved on these Phos-tag gels because of the relatively low protein loading and long run time needed to resolve phosphorylated AHL10 in total protein extracts. E. Effect of HAI1 on *in vivo* phosphorylation status of AHL10 in the −1.2 MPa treatment analyzed by introducing *AHL10*_*promoter*_::*AHL10-YFP/ahl10-1* into *hai1-2* and *35S:FLAG-HAI1* backgrounds. Aliquots of the same samples were run on Phos-tag (top) and SDS-PAGE (middle) gels and band intensities of AHL10-YFP quantified (25 μg protein loaded per lane for both gels). The SDS-PAGE blot was stripped and re-probed to detect HSC70 as a loading control. “C” indicates samples from the unstressed control while “S” indicates stress treatment (−1.2 MPa, 96 h). The experiment was repeated with consistent results. Numbers below each lane indicate relative quantitation of band intensities from the same regions of interest indicated in D.

To test which of these multiple bands of phosphorylated AHL10 were associated with S313 or S314, we immunoprecipitated non-mutated AHL10 (N.M.), AHL10^S313A^ and AHL10^S314A^ from *3S:YFP-AHL10/ahl10-1* (SI Appendix Fig. S8) after −1.2 MPa stress treatment. These proteins were then compared to C.I.P. dephosphorylated AHL10 (Fig. 2B). This showed that the most abundant forms of highly phosphorylated AHL10 (bands 1 and 2 in Fig.2B) were either absent or shifted down in AHL10^S313A^and AHL10^S314A^ while the abundance of a less phosphorylated form (band 3) increased. The slow migration of bands 1 and 2 on Phos-tag gels was consistent with multiple phosphorylation. Likely these additional phosphorylations were at the S297, T311, S317 sites observed in other phosphoproteomic studies; however, phosphorylation at other yet-to-be-identified sites cannot be ruled out. This loss of highly phosphorylated AHL10 in AHL10^S313A^ and AHL10^S314A^ indicated that phosphorylation at these sites may in turn influence phosphorylation of other sites on AHL10. Since bands 1 and 2 were also removed or downshifted by HAI1 treatment (Fig. 2A), these result suggested that HAI1 dephosphorylation of AHL10 could involve S313 and S314.

To further test the effect of HAI1 on AHL10 phosphorylation, AHL10^S313A^ and AHL10^S314A^, as well as the non-mutated (N.M.) AHL10, were used as substrates for *in vitro* dephosphorylation (Fig. 2C). Here again, the most highly phosphorylated bands were eliminated by HAI1 treatment. The downshifted band that accumulated in AHL10 S314A (band “a” in Fig. 2C) could also be seen in AHL10 N.M. after HAI1 treatment, consistent with HAI1 dephosporylation of S314. Interestingly though, AHL10^S313A^ and AHL10^S314A^ incubated with HAI1 were missing band “a” and had decreased intensity of band “b” (Fig. 2C). The most likely explanation for these complex patterns is that the different phosphorylation sites of AHL10 are not independent of one another and that blocking phosphorylation at AHL10^S313A^ and AHL10^S314A^ influences phosphorylation at other sites (for example, phosphorylation at S313 or S314 could potentiate phosphorylation at additional sites or vice versa). At least some of these additional sites can be dephosphorylated by HAI1. While further experiments will be needed to work out mechanisms of AHL10 sequential or cooperative phosphorylation, our data confirm that AHL10 is phosphorylated at multiple sites and that HAI1 can directly dephosphorylate specific sites on AHL10.

To analyze the effect of low ψ_w_ on AHL10 phosphorylation *in planta*, total protein extracts from control and low ψ_w_ treatments were separated on Phos-tag gels (Fig 2D and E). To maintain Phos-tag gel resolution, a limited amount of protein was loaded (25 mg, compared to 100 mg of total protein used for AHL10 immunoprecipitation). Thus only the most abundant bands of phosphorylated AHL10 could be detected. In the moderate severity low ψ_w_ treatment used in growth and gene expression assays (−0.7 MPa, see below), there was a stress induced increase in both phosphorylated AHL10 and total AHL10 protein (compare Phos-tag and SDS-PAGE in Fig. 2D). The two major phosphorylated bands observed were dependent on S313 and S314 phosphorylation as both bands were absent or shifted in the AHL10^S313A^ and AHL10^S314A^ (Fig. 2D). This was consistent with the results shown in Fig 2B and similar results were observed with AHL10 immunoprecipitated from seedlings at −0.7 MPa (SI Appendix Fig. S8D). In the −1.2 MPa treatment used for phosphoproteomics, there was strong increase in phosphorylated AHL10 and moderate increase in AHL10 protein level (Fig. 2E). This indicated a stress-induced increase in AHL10 phosphorylation stoichometry at −1.2 MPa. Note that because the bands of phosphorylated AHL10 were more intense at −1.2 MPa (Fig. 2E), a shorter exposure was used compared to the −0.7 MPa immunoblot (Fig. 2D) to avoid saturation and thus the phosphorylated bands in the control were not as clearly visible in Fig. 2E compared to Fig. 2D.

Further increased abundance of phosphorylated AHL10 was seen in *hai1-2* compared to wild type at −1.2 MPa (Fig. 2E). This was consistent with HAI1 regulation of AHL10 phosphorylation and consistent with our phosphoproteomics data. Conversely, HAI1 ectopic expression (*35S:HAI1*) caused the amount of phosphorylated AHL10 to decrease in the control but had no additional effect during −1.2 MPa stress, presumably because endogenous HAI1 was already highly expressed. Together these results indicated that HAI1 restricted the level of phosphorylated AHL10. Despite the action of HAI1, abundance of phosphorylated AHL10 increased during low ψ_w_ stress. This was partially because of increased AHL10 protein level and also likely because of increased kinase activity to phosphorylate AHL10. As S314 is followed by a proline (SP phosphorylation motif), and the equivalent site on AHL13 has been identified as a putative MPK target (9), it is possible that stress-activated MPK activity is responsible for the increased AHL10 phosphorylation at low ψ_w_. Interestingly, *hai1-2* and *35S:HAI1* had decreased AHL10 protein abundance in the unstressed control (Fig. 2D and E; SI Appendix Fig. S8D). Thus it is also possible that HAI1 influences AHL10 protein stability, either through dephosphorylation or via other indirect mechanisms.

We focused further functional analysis (below) on AHL10 S313 and S314 as these were the phosphorylation sites most frequently identified in phosphoproteomic studies, including our own. Also, multiple sets of the Phos-tag gel analyses indicated that S313 and S314 were the dominant sites of AHL10 phosphorylation because they were required for the most abundant forms of phosphorylated AHL10, either by being the most heavily phosphorylated sites themselves and also because they may be required for phosphorylation at other sites, including additional sites dephosphorylated by HAI1.

### AHL10 phosphorylation at S314 is critical for suppression of growth during moderate severity low ψ_w_ stress

The effect of HAI1 on AHL10 phosphorylation and increased protein level of AHL10 during low ψ_w_ suggested that AHL10 may also be involved in low ψ_w_ response. Consistent with this, *ahl10-1* had nearly forty percent higher rosette weight than wild type after an extended period of moderate severity soil drying (Fig. 3A, B). A similar enhanced growth maintenance of *ahl10-1* was seen in plate-based assays where seedlings were transferred to fresh control plates or to moderate severity low ψ_w_ plates (−0.7 MPa) and root elongation measured over the next five days in the control and 10 days in the low ψ_w_ stress treatment (Fig. 3C, D; wild type root length and dry weights as well as fresh weight of all genotypes at the time of transfer are shown in SI Appendix Fig. S9). The increased seedling weight and root elongation of *ahl10-1* could be complemented by *AHL10*_*promotor*_:*AHL10-YFP* and *35S:YFP-*AHL10 (Fig. 3C and D; *35S:YFP-AHL10* is labeled “N.M.” for Not Mutated). *hai1-2* also had increased seedling weight and root elongation at low ψ_w_. The same pattern was observed when seedlings were transferred to the −1.2 MPa stress treatment (SI Appendix Fig. S10A-C). Although the −1.2 MPa treatment restricted growth to a low level, it was not lethal and seedlings subjected to −1.2 MPa for ten days rapidly recovered when returned to the unstressed control media (SI Appendix Fig. S10D).

**Figure 3:**
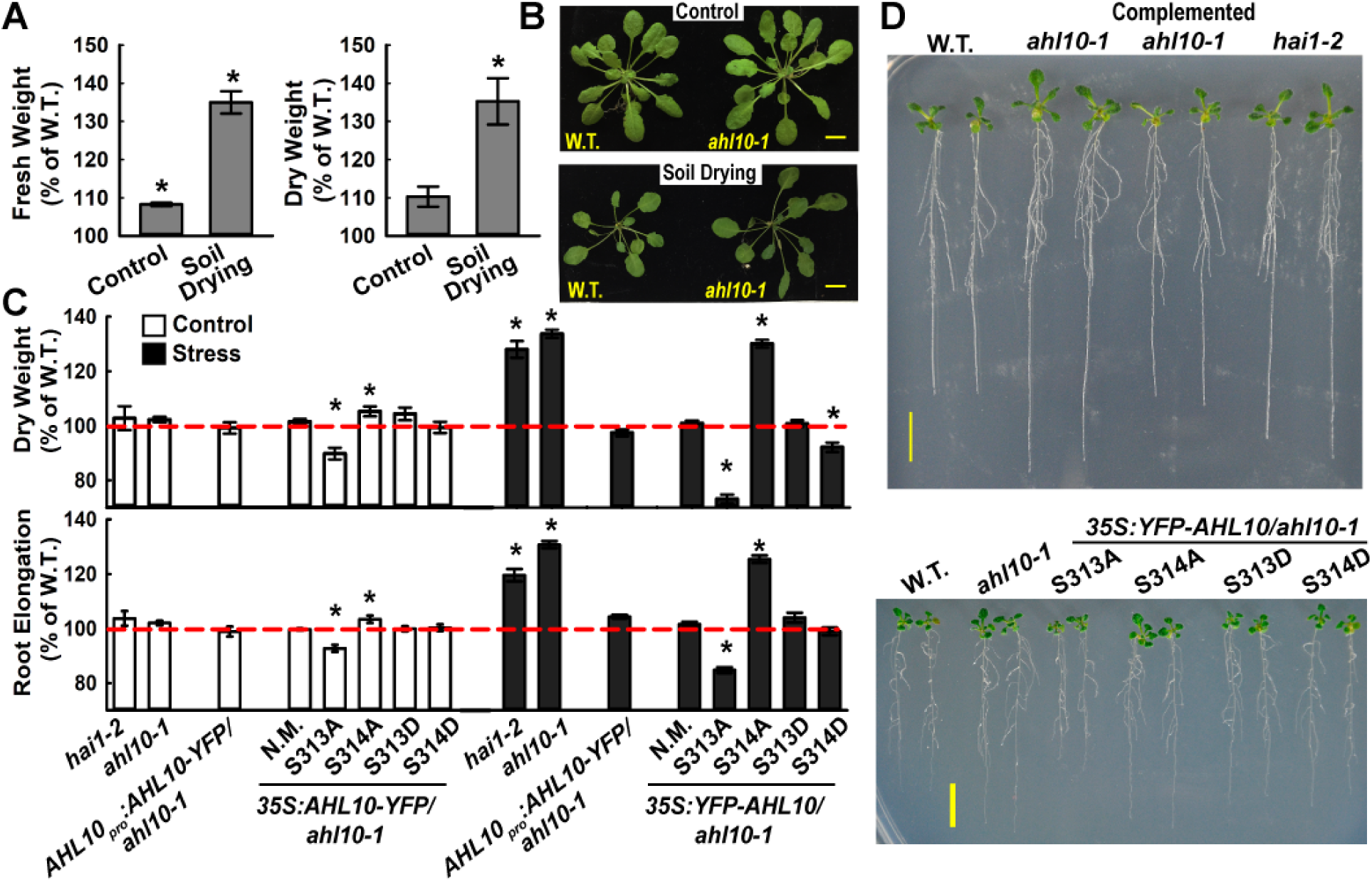
AHL10 phosphorylation at S314 is critical for growth suppression during low ψw stress. A. Relative Rosette fresh weight and dry weight of *ahl10-1* compared to Col-0 wild type in control or soil drying treatments (mean ± SE, n = 14-16 combined from three independent experiments. Asterisk (*) indicates significant difference (P ≤ 0.05) compared to wild type (100%) by one-sample t test). Wild type mean rosette fresh weight across all three experiments was 256.2 ± 23.8 mg and 72.3 ± 4.2 mg in the well-watered control and soil drying treatments, respectively. Wild type mean dry weight was 18.3 ±1.8 mg and 7.4 ± 0.4 mg in the control and soil drying treatments, respectively. B. Representative rosettes of the wild type and *ahl10-1* in control and soil drying treatments. Scale bars = 1 cm. C. Root elongation and dry weight of seedlings under control and low ψ_w_ stress (−0.7 MPa) conditions. Data are relative to the Col-0 wild type (mean ± SE, n = 30 to 45, combined from three independent experiments). Asterisk (*) indicates significant difference compared with the wild type by one-sample T-test (P ≤ 0.05). Dashed red line indicates the wild type level (100%). Seedling weights and root elongation of Col-0 wild type used for normalization are shown in SI Appendix Fig S9. The mean dry weight of wild type seedlings was 0.64 ± 0.01 mg and 0.67 ± 0.02 mg in the control and −0.7 MPa stress treatment, respectively. The mean root elongation of wild type was 63.9 ± 0.26 mm and 22.5 ± 0.3 mm in the control and −0.7 MPa treatments, respectively (these data are also shown in SI Appendix Fig S9D). Note that three independent transgenic lines were analyzed for each construct and combined data of all three is shown. For the *35S:YFP-AHL10/ahl10*, N.M. indicates Non Mutated (ie: wild type) AHL10. D. Representative seedlings of Col-0 wild type (W.T.), *ahl10-1, AHL10*_*promoter*_:*AHL10-YFP/ahl10-1* (Complemented *ahl10-1*) and *hai1-2* as well as *ahl10-1* complemented with, phosphonull AHL10 (S313A, S314A) or phosphomimic AHL10 (S313D, S314D) all expressed under control of the *35S* promoter (*35S:YFP-AHL10/ahl10-1*). Five-day-old seedlings were transferred to −0.7 MPa and pictures taken 10 days after transfer when the quantitation of root elongation and seedling weight was performed. Scale bars = 1 cm.

To test the functional importance of AHL10 phosphorylation, phosphonull (S313A or S314A) and phosphomimic (S313D or S314D) AHL10 were expressed in the *ahl10-1* background. AHL10^S314A^ was unable to complement *ahl10-1* while AHL10^S314D^ and AHL10^S313D^ did complement (Fig. 3C, D). Interestingly, AHL10^S313A^ suppressed growth below the wild type level in both control and stress treatments. The same pattern was observed in the −1.2 MPa treatment (SI Appendix Fig. S10B, C). These data show that AHL10 function in growth regulation is dependent on S314 phosphorylation. The greater suppression of growth in AHL10^S313A^ indicates that S313 may act as a “decoy” site such that either S313 or S314 can be phosphorylated by the kinase(s) acting on AHL10. When S313 phosphorylation is blocked more S314 phosphorylation occurs leading to increased AHL10 function in growth suppression.However, other mechanisms, perhaps involving coordinated phosphorylation at additional sites on AHL10 as mentioned above, cannot be ruled out. Note that three independent transgenic lines were used for each AHL10 construct and each line had indistinguishable growth phenotypes despite some difference in AHL10 protein level (SI Appendix Fig. S8B). Thus, the combined data of all three lines for each construct is shown in Fig. 3C. All of the AHL10 phosphomimic and null proteins were localized in the nucleus; however, we noted that localization of AHL10^S313D^ and AHL10^S314D^ was less diffuse and partially clustered into foci (SI Appendix Fig. S8C).

### AHL10 regulates the expression of developmental and hormone-associated genes during drought acclimation

To learn how AHL10 could influence growth, RNAseq was conducted for wild type and *ahl10-1* under unstressed conditions or moderate severity low ψ_w_ stress (−0.7 MPa, 96 h) where the growth maintenance phenotype of *ahl10-1* was apparent. In wild type 2212 genes were up-regulated and 2766 down regulated at low ψ_w_ (Dataset S6, S7; fold change in expression >1.25, adjusted P ≤ 0.05). *HAI1* was strongly upregulated by low ψ_w_ (Dataset S6) in agreement with previous data (15). In the unstressed control, *ahl10-1* had little effect on gene expression as only 10 genes had higher expression and 7 genes (excluding *AHL10* itself) had reduced expression compared to wild type (Dataset S8). There was more effect of *ahl10-1* at low ψ_w_ where 19 genes were increased and 41 genes decreased compared to wild type (Dataset S9). The majority of genes differentially expressed in *ahl10-1* under stress were also differentially expressed in wild type at low ψ_w_ compared to the unstressed control. However, there was no concordance between the effect of stress in wild type (increased or decreased expression) and the direction of the *ahl10-1* effect (Fig. 4A). Only seven genes were differentially expressed in *ahl10-1* in both control and low ψ_w_ treatments (Fig. 4A, inset). Six of these were transposons or “other RNA” rather than protein coding genes.

**Figure 4:**
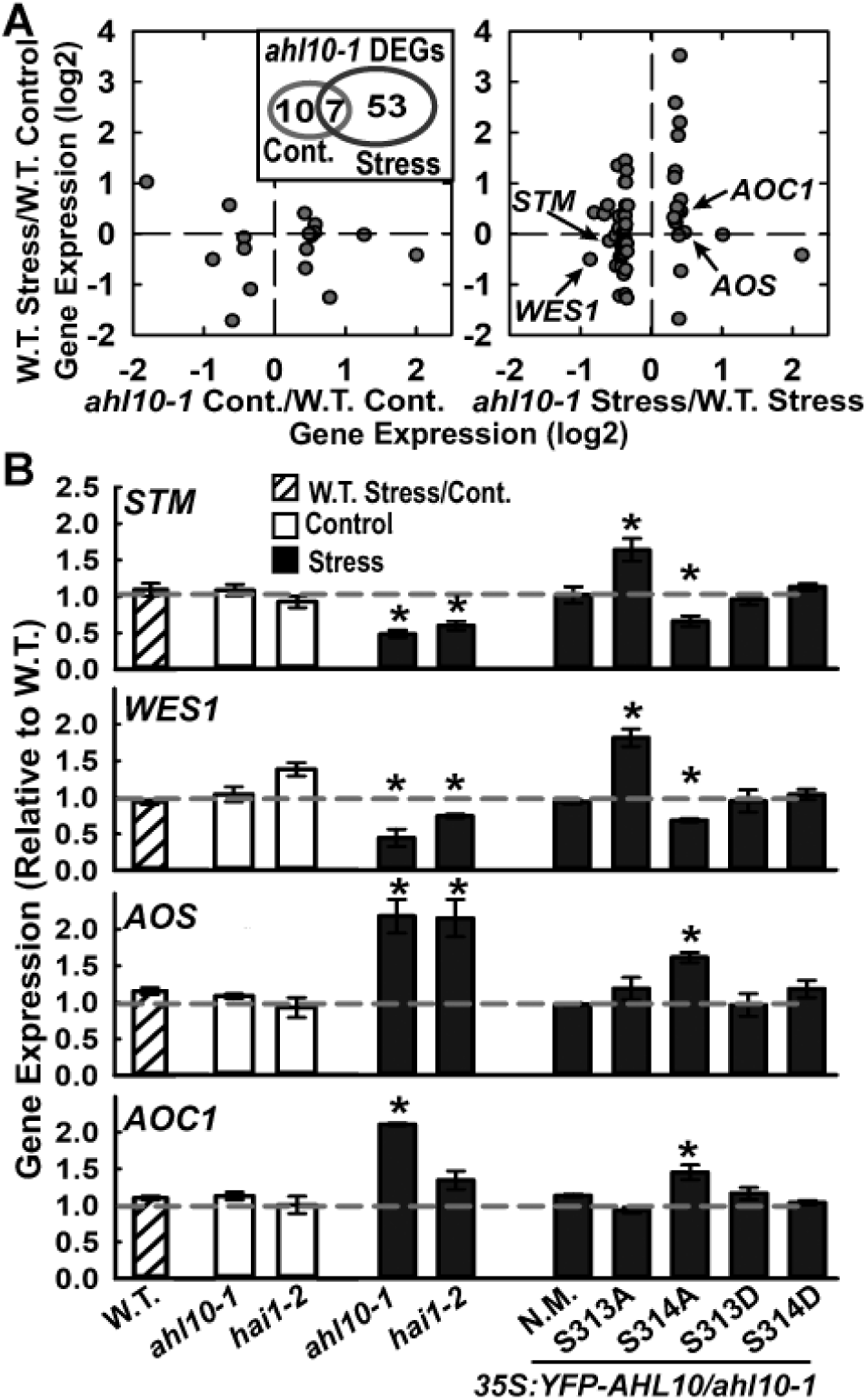
AHL10 regulation of development and hormone-related genes during low ψ_w_ stress also depends upon S314 phosphorylation. A. Differentially expressed genes (DEGs) in *ahl10-1* discovered by RNAseq. Genes with differential expression in *ahl10-1* under either control or stress (−0.7 MPa, 96 h) plotted versus their expression in wild type stress versus wild type control. Inset shows the overlap between genes with differential expression in *ahl10-1* in control or stress treatments. B. QPCR assay of selected genes in stress (−0.7 MPa, 96 h) versus control for wild type as well as assay of *ahl10-1, hai1-2* and AHL10 phosphomimic and null complementation lines. Data are shown as expression relative to wild type and are means ± SE, (n = 3) combined from three independent experiments. Asterisk (*) indicates significant difference compared to wild type in the same treatment (or wild type stress versus control) by one-sample T-test (P ≤ 0.05). Gray dashed line indicates the wild type level (set to 1).

The genes differentially expressed in *ahl10-1* at low ψ_w_ were consistent with altered growth regulation. These included *Shootmeristemless* (*STM*), a transcription factor required for meristem maintenance (39, 40) and two auxin-amido synthase genes (*WES1* and *DFL1*) that affect growth by controlling the level of active auxin (41–43). *WES1* is also known to be regulated by stress and ABA (41). Genes with increased expression in *ahl10-1* at low ψ_w_ included *Root Meristem Growth Factor 9* (*RGF9*), which is one of a family of genes that affect proliferation of transient amplifying cells in the root meristem (44), as well as the JA synthesis genes *Allene Oxide Synthase* (*AOS*) and *Allene Oxide Cyclase* (*AOC1*) and JA-responsive genes *JR2* and *VSP2* (45, 46). Many of these genes, for example *STM* and *RGF9*, have tissue specific expression. Thus their change in expression in meristems or other specific tissues may be more dramatic more than that seen in the whole seedling data shown here.

Quantitative RT-PCR validated that *STM* and *WES1* (Fig. 4B) as well as other genes (SI Appendix Fig. S11A) had decreased expression in *ahl10-1* at low ψw. Similarly, *AOS, AOC1* (Fig. 4B) and other genes (SI Appendix Fig. S11B) were confirmed to have increased expression in *ahl10-1* at low ψ_w_. Consistent with the growth assays, AHL10^S314A^ was unable to complement the altered gene expression of *ahl10-1* while non-mutated (N.M.) AHL10, AHL10^S313D^ and AHL10^S314D^could complement (Fig. 4B). For *STM* and *WES1* we again observed that AHL10^S313A^ acted as a hyper-functional allele having the opposite effect as *ahl10-1* (Fig. 4B). Also consistent with the growth assays, *hai1-2* had similar effect on gene expression at low ψw compared to *ahl10-1* (Fig. 4B).

### AHL10 S314 phosphorylation does not affect AHL10 self-interaction but is required for AHL10 complexes to form nuclear foci

The above results raised the question of how phosphorylation affects AHL10 protein function. Structural modeling confirmed that AHL10 S313 and S314 are in a loop region not immediately adjacent to the two AT-hook DNA binding domains or the PPC/DUF296 domain involved in AHL trimer formation (SI Appendix Fig. S12). This variable C-terminal portion of AHLs is thought to be involved in interaction of AHL complexes with other transcriptional regulators (47). AHL10 S313 and S314 phosphomimic and phosphonull variants had no effect on AHL10 self-interaction in rBiFC assays (Fig. 5A, rBiFC representative images are in SI Appendix Fig. S13). Strikingly, for non-mutated AHL10 (N.M.), phosphomimic (S313D, S314D) and AHL10^S313A^, the rBiFC fluorescence from AHL10 complexes could often be seen as foci within the nucleus (Fig. 5B). The portion of nuclei with AHL10 foci increased under stress and was dramatically increased by the S313A phosphonull mutation (Fig. 5C). In contrast, AHL10^S314A^ did not form foci and remained dispersed in the nucleoplasm (Fig. 5B, C). These data, along with reports that AT-hook proteins can associate with the nuclear matrix (48, 49), raise the possibility that interaction with the nuclear matrix or recruitment of AHL10 complexes to matrix attachment regions is dependent upon S314 phosphorylation.

**Figure 5:**
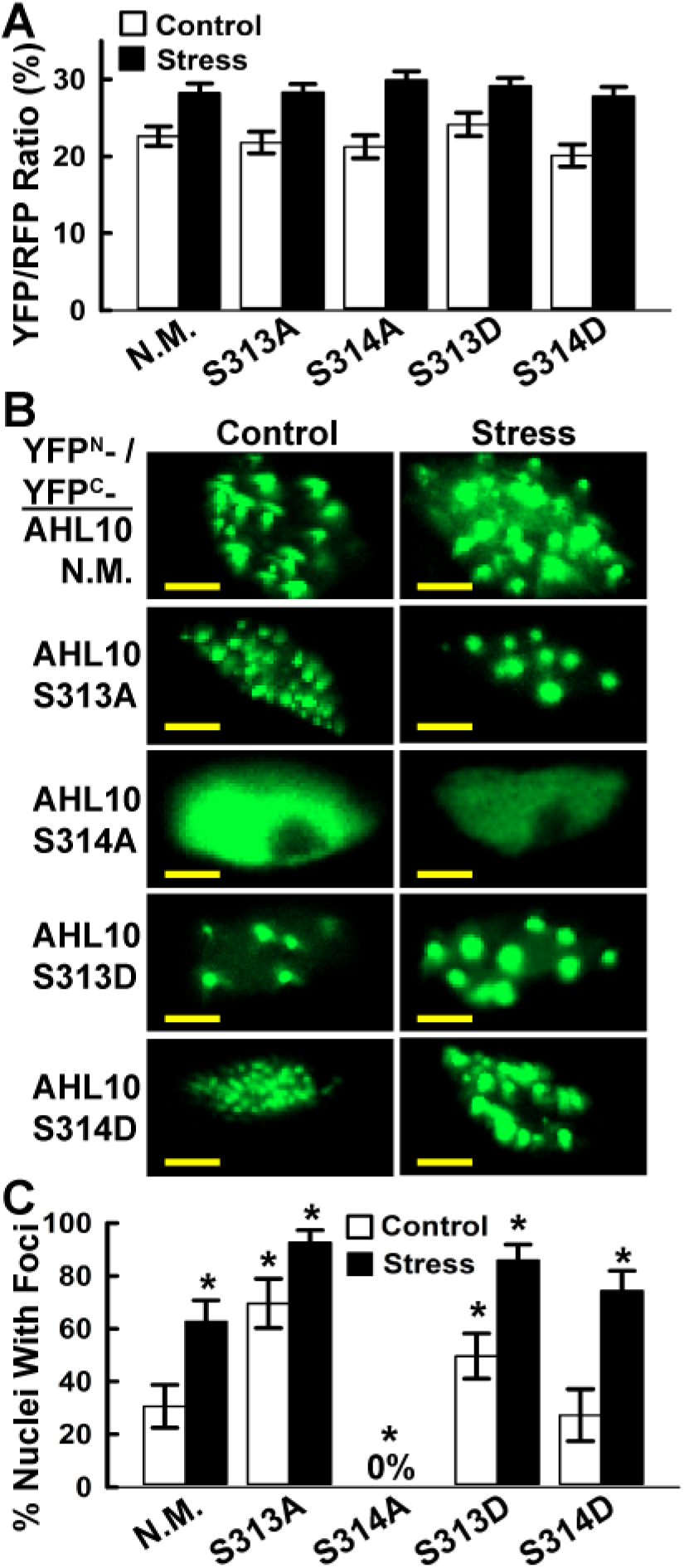
Ratiometric BiFC (rBiFC) analysis of self-interaction and nuclear foci localization of phosphomimic and null AHL10. A. Quantification of relative self-interaction intensity for phosphomimic and phosphonull AHL10. Data are means ± SE, (n = 10 to 20) combined from two independent experiments. None of the phosphomimic or phosphonull constructs differed significantly from wild type in either stress or control treatments (T-test P ≤ 0.05). Images of AHL10 self-interaction rBiFC assays are shown in SI Appendix Fig. S13. B. Representative images of nuclear foci localization of AHL10 self-interaction complexes observed in rBiFC assays. For non-mutated wild type AHL10 (N.M.) and phosphomimic AHL10^S313D^ and AHL10^S314D^, representative images of nuclei without foci are shown in SI Appendix Fig. S13B. Note that nuclei with foci were never observed for AHL10^S313A^. Scale bars indicate 5 μm. C. Portion of nuclei with AHL10 foci. Individual nuclei (80-130 for each construct and treatment) combined from two independent experiments were counted. Error bars indicate 95% confidence intervals and asterisks indicate significant difference compared to wild type in the same treatment (or difference of wild type stress versus control) based on Fisher’s Exact test (P ≤ 0.05).

## Discussion

We identified a diverse set of proteins with altered phosphopeptide abundance in *hai1-2* compared to wild type. Given the strong effect of HAI1 on drought-related phenotypes, many of these HAI1-affected phosphoproteins may have previously unknown roles in drought response. The phosphorylation sites we identified likely represent a combination of sites directly dephosphorylated by HAI1 and sites phosphorylated by HAI1-regulated kinases (or both).AHL10 is a good example as the S314 phosphorylation site contained a Ser-Pro phosphorylation motif that is typically targeted by MPKs and the analogous phosphoserine site on AHL13 was identified as a putative MPK target (9). Thus, even though HAI1 could dephosphorylate AHL10 *in vitro*, AHL10 phosphorylation *in vivo* may also be affected by HAI1 dephosphorylation of MPKs (or other kinases) as well as activation of MPK activity by environmental stimuli (see model in SI Appendix Fig. S14). The growth assays, gene expression analysis and localization of AHL10 complexes together demonstrated that S314 phosphorylation was a critical determinant of AHL10 function. These assays also indicated that S313 phosphorylation had an opposing effect, possibly by affecting the level of S314 phosphorylation. Consistent with these data, Phos-tag gel analysis showed that the major bands of phosphorylated AHL10 that increased in abundance during low ψ_w_ were absent or shifted in AHL10^S313A^ and AHL10^S314A^. These bands were also eliminated or shifted by in vitro dephosphorylation with HAI1. While these data point to S313 and S314 as the critical phosphorylation sites for AHL10 function, we cannot exclude the involvement of other phosphorylation sites. Search of the PhosPhat database found five experimentally validated phosphorylation sites in a 20 amino acid stretch near the C-terminus of AHL10 (S297, T311, S313, S314, S317). It is possible that part of the importance of S313 and S314 phosphorylation is to influence phosphorylation at additional nearby sites. The complex pattern of phosphorylation changes revealed by HAI1 *in vitro* dephosphorylation of AHL10^S313A^ and AHL10^S314A^ suggest that this may be true. Clusters of phosphorylation sites are observed on many proteins (50) and numerous examples of cooperative phosphorylation regulating protein function can be found among well studied metazoan proteins.

AHL10 acted as a negative regulator to restrain growth during low ψ_w_ stress. In some ways this effect of AHL10 was consistent with Clade A AHL (AHL15-AHL27) suppression of hypocotyl elongation (47, 51, 52). However, our results differ in that AHL10 primarily affected growth at low ψ_w_ and in that AHL10 is a member of the less studied Clade B AHLs (AHL1-AHL14). The only other study of AHL10 that we are aware of found AHL10 to be involved in recruitment of the heterochromatin mark Histone 3 Lysine 9 dimethylation (H3K9me2) to AT-rich transposable elements (53). We observed mis-regulated expression of several transposons in *ahl10-1*. However, these were a minority of the differentially expressed genes, especially at low ψ_w_. Whether or not histone modification or other epigenetic mechanisms are involved in AHL10-mediated induction (for example *STM, WES1, DFL1*) or repression (for example *AOS, AOC1*) of gene expression is of interest for future research. Interestingly, AHL10 and AHL13, with phosphorylation at S313/S314 or the analogous AHL13 site, were the only AHLs identified in phosphoproteomic analysis of chromatin associated proteins (54). When considered along with other data presented here, this suggests that AHL10 and AHL13 may differ in function and phospho-regulation compared to other AHLs. The only previous report of AHL involvement in drought response found that overexpression of a rice AHL, which the authors designated as *OsAHL1*, led to enhanced survival of dehydration, salt stress or chilling stress (55). While this may at first seem inconsistent with our results, it is important to note that they measured survival of severe stress while we measured the response to low ψ_w_ treatments that inhibited growth but were not lethal. It has been well noted that the mechanisms underlying survival of severe stress are distinct from those regulating growth during moderate severity drought stress (56).

Altered expression of developmental regulators *STM* and *RGF9,* auxin metabolism genes *WES1* and *DFL1* and JA metabolism genes *AOS, AOC,* as well as other AHL10-affected genes identified by RNAseq, was consistent with AHL10 effects on growth at low ψ_w_. Many of these genes have tissue specific expression (for example STM). Thus, tissue specific analysis of meristematic and growing regions could find additional AHL10-regulated genes important for growth regulation or reveal much stronger localized changes in expression of the AHL10-regulated genes we identified. Mine et al. (19) demonstrated that JA-induction of *HAI1* expression was involved in HAI1 suppression of MPK3 and MPK6 activation during effector triggered immunity. Our finding that HAI1-AHL10 signaling regulated JA metabolism (*AOS* and *AOC1*) and JA responsive (*JR2, VSP2*) genes suggests that HAI1 also has a feedback effect on JA. This result is also consistent with other reports of cross-regulation between JA and ABA metabolism (57). In our previous characterization of HAI1, we noted that *hai1-2* had altered expression of many defense-related genes and concluded that HAI1 could be involved in trade-offs between pathogen defense and drought response (15). The current data further indicate that HAI1, AHL10 and their target genes are involved in this intersection of stress and defense signaling with growth regulation.

AHLs can associate with AT-rich sequences in matrix attachment regions (48, 49) and the C-terminal portion of AHLs is thought to mediate interaction with transcriptional regulators (47). AHL10 S314 phosphorylation was required for AHL10 complexes to form foci within the nucleoplasm. While the identity and composition of the AHL10 foci is not known, it can be hypothesized that phosphorylation-dependent interaction of AHL10 with nuclear matrix components as well as other transcriptional regulators or epigenetic factors may explain how AHL10 could act as either an inducer or suppressor of gene expression under low ψw.Interestingly, AHL^S313A^ had a hyperactive effect on the formation of nuclear foci and on *STM* and *WES1* expression but not on *AOS1* or *AOC* expression (Fig. 3B). This provides one indication that different mechanisms may be involved in AHL10 up and down-regulation of gene expression. As AHL10 foci were observed in both control and low ψ_w_. treatments, it is possible that the AHL10 foci are involved in repression of transposons (54), which was disrupted under both control and stress conditions in *ahl10-1*. Other, yet to be identified, phosphorylation dependent interactions of AHL10 may be involved in stress-specific effects on gene expression.

Both previous data (15) and RNAseq analysis conducted here (Dataset S6, S7) show that *HAI1* and *HAI2* are among the genes most strongly induced during low ψ_w_ acclimation. Along with previous physiological results (15, 19, 58), such data indicate the prominent and distinct role that the HAI PP2Cs play in plant responses to the environment. As illustrated by our characterization of AHL10, the *hai1-2* phosphoproteomics data provide a basis to better understand the phosphatase side of the kinase-phosphatase interplay that is a central feature of stress signaling. More broadly the data indicate roles for AHL10 and HAI1 in balancing growth versus stress and defense responses. AHL10-mediated growth suppression under moderate stress severity can be an adaptive response to reduce water use. Agronomically, such a strategy may be overly conservative and disrupting it could allow greater plant productivity in certain environments.

## Materials and Methods

Low ψ_w_. stress was imposed using PEG-infused agar plates or controlled soil drying as previously described (59, 11). For phosphoproteomics, seedlings were transferred to unstressed control or −1.2 MPa low ψ_w_. stress and samples collected 96 h after transfer. Phosphoproteomics sample processing, phosphopeptide enrichment, iTRAQ labeling, and data analysis for phosphopeptide quantitation are as described previously (11) and are fully detailed in the SI Materials and Methods. Construction of transgenic plants, phenotypic assays and protein analyses were performed generally as previously described (11) with further details and modifications described in SI Materials and Methods. For expression of recombinant HAI1, an N-terminal truncation lacking amino acids 1-103, similar to the truncated HAI1 previously used for in vitro assays (30) was expressed in *E.coli* and purified as described in SI Materials and Methods. Immunoprecipitation of YFP-tagged AHL10, in vitro dephosphorylation assays and Phos-tag gel analysis are described in SI Appendix Materials and Methods. Transient expression for BiFC assays in Arabidopsis seedlings used a protocol modified from (60) and ratiometric BiFC vector system (61) or traditional BiFC vectors as described previously (11) and in SI Materials and Methods. Phosphoproteomics data have been submitted to the Proteome Exchange Consortium with the data set identifier PXD004869 and RNAseq data can be accessed in the Gene Expression Omnibus (GEO) repository with accession number GSE112368.

## Supporting information

## Acknowledgements

We thank T.Z. Chang and S.-S. Huang for laboratory assistance, M.-J. Fang and J.-Y Huang for microscopy assistance and M.-Y. Lu (High Throughput Sequencing Core, Biodiversity Research Center, Academia Sinica) for RNAseq service and T.E. Juenger (University of Texas-Austin) for statistical advice. This work was supported by Taiwan Ministry of Science and Technology Grant 103-2314-B-001-003 (to P.E.V.).

## Supplemental Materials Materials and Methods

Figures S1-S14

Supplemental Datasets S1-S10

## References

1 Finkelstein R (2013) Abscisic Acid Synthesis and Response. The Arabidopsis Book:e0166.

2 Verslues PE (2016) ABA and cytokinins: challenge and opportunity for plant stress research. Plant Molec Biol 91(6):629–640.

3 Cutler SR, Rodriguez PL, Finkelstein RR, & Abrams SR (2010) Abscisic Acid: Emergence of a Core Signaling Network. Ann Rev Plant Biol 61:651–679.

4 Raghavendra AS, Gonugunta VK, Christmann A, & Grill E (2010) ABA perception and signalling. Trends Plant Sci 15(7):395–401.

5 de Zelicourt A, Colcombet J, & Hirt H (2016) The Role of MAPK Modules and ABA during Abiotic Stress Signaling. Trends Plant Sci 21(8):677–685.

6 Umezawa T, et al. (2013) Genetics and Phosphoproteomics Reveal a Protein Phosphorylation Network in the Abscisic Acid Signaling Pathway in Arabidopsis thaliana. Sci. Signal. 6(270):270.

7 Wang PC, et al. (2013) Quantitative phosphoproteomics identifies SnRK2 protein kinase substrates and reveals the effectors of abscisic acid action. Proc Natl Acad Sci USA 110(27):11205–11210.

8 Minkoff BB, Stecker KE, & Sussman MR (2015) Rapid Phosphoproteomic Effects of Abscisic Acid (ABA) on Wild-Type and ABA Receptor-Deficient A-thaliana Mutants. Molec Cell Prot 14(5):1169–1182.

9 Hoehenwarter W, et al. (2013) Identification of Novel in vivo MAP Kinase Substrates in Arabidopsis thaliana Through Use of Tandem Metal Oxide Affinity Chromatography. Molec Cell Prot 12(2):369–380.

10 Sörensson C, et al. (2012) Determination of primary sequence specificity of Arabidopsis MAPKs MPK3 and MPK6 leads to identification of new substrates. Biochem J 446(2):271–278.

11 Bhaskara GB, Wen TN, Nguyen TT, & Verslues PE (2017) Protein Phosphatase 2Cs and Microtubule-Associated Stress Protein 1 Control Microtubule Stability, Plant Growth, and Drought Response. Plant Cell 29(1):169–191.

12 Leung J, et al. (1994) Arabidopsis ABA response gene ABI1: features of a calcium-modulated protein phosphatase. Science 264(5164):1448.

13 Meyer K, Leube MP, & Grill E (1994) A protein phosphatase 2C involved in ABA signal transduction in Arabidopsis thaliana. Science 264(5164):1452.

14 Saez A, et al. (2004) Gain-of-function and loss-of-function phenotypes of the protein phosphatase 2C HAB1 reveal its role as a negative regulator of abscisic acid signalling. Plant J 37(3):354–369.

15 Bhaskara GB, Nguyen TT, & Verslues PE (2012) Unique Drought Resistance Functions of the Highly ABA-Induced Clade A Protein Phosphatase 2Cs. Plant Physiol 160(1):379–395.

16 Yoshida T, et al. (2006) ABA-hypersensitive germination3 encodes a protein phosphatase 2C (AtPP2CA) that strongly regulates abscisic acid signaling during germination among Arabidopsis protein phosphatase 2Cs. Plant Physiol 140(1):115–126.

17 Fujita Y, et al. (2009) Three SnRK2 Protein Kinases are the Main Positive Regulators of Abscisic Acid Signaling in Response to Water Stress in Arabidopsis. Plant Cell Physiol 50(12):2123–2132.

18 Wang K, et al. (2018) EAR1 negatively regulates ABA signaling by enhancing 2C protein phosphatase activity. Plant Cell 30(4):815–834

19 Mine A, et al. (2017) Pathogen exploitation of an abscisic acid-and jasmonate-inducible MAPK phosphatase and its interception by Arabidopsis immunity. Proc Natl Acad Sci USA 114(28):7456–7461.

20 Iyer-Pascuzzi Anjali S, et al. (2011) Cell Identity Regulators Link Development and Stress Responses in the Arabidopsis Root. Develop Cell 21(4):770–782.

21 Bhaskara GB, Nguyen TT, Yang TH, & Verslues PE (2017) Comparative Analysis of Phosphoproteome Remodeling After Short Term Water Stress and ABA Treatments versus Longer Term Water Stress Acclimation. Front Plant Sci 8: 523.

22 Vlad F, Turk BE, Peynot P, Leung J, & Merlot S (2008) A versatile strategy to define the phosphorylation preferences of plant protein kinases and screen for putative substrates. Plant J 55(1):104–117.

23 Amanchy R, et al. (2007) A curated compendium of phosphorylation motifs. Nat Biotech 25(3):285–286.

24 Takahashi Y, Ebisu Y, & Shimazaki K (2017) Reconstitution of Abscisic Acid Signaling from the Receptor to DNA via bHLH Transcription Factors. Plant Physiol 174(2):815– 822.

25 Merkouropoulos G, Andreasson E, Hess D, Boller T, & Peck SC (2008) An Arabidopsis protein phosphorylated in response to microbial elicitation, AtPHOS32, is a substrate of MAP kinases 3 and 6. J Biol Chem 283(16):10493–10499.

26 de la Fuente van Bentem S, et al. (2008) Site-Specific Phosphorylation Profiling of Arabidopsis Proteins by Mass Spectrometry and Peptide Chip Analysis. J Prot Res 7(6):2458–2470.

27 Huck NV, et al. (2017) Combined 15N-Labeling and TandemMOAC Quantifies Phosphorylation of MAP Kinase Substrates Downstream of MKK7 in Arabidopsis. Front Plant Sci 8(2050):2050.

28 Zhao SS, et al. (2016) CASEIN KINASE1-LIKE PROTEIN2 Regulates Actin Filament Stability and Stomatal Closure via Phosphorylation of Actin Depolymerizing Factor. Plant Cell 28(6):1422–1439.

29 Cui Y, et al. (2014) Arabidopsis casein kinase 1-like 2 involved in abscisic acid signal transduction pathways. J Plant Interact 9(1):19–25.

30 Antoni R, et al. (2012) Selective inhibition of clade A phosphatases type 2C by PYR/PYL/RCAR abscisic acid receptors. Plant Physiol 158(2):970–980

31 Ufer G, Gertzmann A, Gasulla F, Röhrig H, & Bartels D (2017) Identification and characterization of the phosphatidic acid-binding A. thaliana phosphoprotein PLDrp1 that is regulated by PLDα1 in a stress-dependent manner. Plant J 92(2):276–290.

32 Choudhary MK, Nomura Y, Wang L, Nakagami H, & Somers DE (2015) Quantitative Circadian Phosphoproteomic Analysis of Arabidopsis Reveals Extensive Clock Control of Key Components in Physiological, Metabolic, and Signaling Pathways. Molec & Cell Proteomics 14(8):2243–2260.

33 Lin L-L, et al. (2015) Integrating Phosphoproteomics and Bioinformatics to Study Brassinosteroid-Regulated Phosphorylation Dynamics in Arabidopsis. BMC Genomics 16(1):533.

34 Nakagami H, et al. (2010) Large-Scale Comparative Phosphoproteomics Identifies Conserved Phosphorylation Sites in Plants. Plant Physiol 153(3):1161–1174.

35 Reiland S, et al. (2009) Large-Scale Arabidopsis Phosphoproteome Profiling Reveals Novel Chloroplast Kinase Substrates and Phosphorylation Networks. Plant Physiol 150(2):889–903.

36 Roitinger E, et al. (2015) Quantitative Phosphoproteomics of the Ataxia Telangiectasia-Mutated (ATM) and Ataxia Telangiectasia-Mutated and Rad3-related (ATR) Dependent DNA Damage Response in Arabidopsis thaliana. Molec & Cell Proteomics 14(3):556– 571.

37 Wang X, et al. (2013) A large-scale protein phosphorylation analysis reveals novel phosphorylation motifs and phosphoregulatory networks in Arabidopsis. J Proteomics 78:486–498.

38 Zhang H, et al. (2013) Quantitative Phosphoproteomics after Auxin-stimulated Lateral Root Induction Identifies an SNX1 Protein Phosphorylation Site Required for Growth. Molec & Cell Proteomics 12(5):1158–1169.

39 Barton MK & Poethig RS (1993) Formation of the shoot apical meristem in Arabidopsis thaliana-An analysis of development in the wild type and in the Shoot Meristemless mutant. Development 119(3):823–831.

40 Landrein B, et al. (2015) Mechanical stress contributes to the expression of the STM homeobox gene in Arabidopsis shoot meristems. Elife 4.

41 Park JE, et al. (2007) GH3-mediated auxin homeostasis links growth regulation with stress adaptation response in Arabidopsis. J Biol Chem 282(13):10036–10046.

42 Nakazawa M, et al. (2001) DFL1, an auxin-responsive GH3 gene homologue, negatively regulates shoot cell elongation and lateral root formation, and positively regulates the light response of hypocotyl length. Plant J 25(2):213–221.

43 Staswick PE, et al. (2005) Characterization of an Arabidopsis enzyme family that conjugates amino acids to indole-3-acetic acid. Plant Cell 17(2):616–627.

44 Matsuzaki Y, Ogawa-Ohnishi M, Mori A, & Matsubayashi Y (2010) Secreted Peptide Signals Required for Maintenance of Root Stem Cell Niche in Arabidopsis. Science 329(5995):1065–1067.

45 Laudert D & Weiler EW (1998) Allene oxide synthase: a major control point in Arabidopsis thaliana octadecanoid signalling. Plant J 15(5):675–684.

46 Leon-Reyes A, et al. (2010) Salicylate-mediated suppression of jasmonate-responsive gene expression in Arabidopsis is targeted downstream of the jasmonate biosynthesis pathway. Planta 232(6):1423–1432.

47 Zhao JF, Favero DS, Peng H, & Neff MM (2013) Arabidopsis thaliana AHL family modulates hypocotyl growth redundantly by interacting with each other via the PPC/DUF296 domain. Proc Natl Acad Sci USA 110(48):E4688–E4697.

48 Fujimoto S, et al. (2004) Identification of a novel plant MAR DNA binding protein localized on chromosomal surfaces. Plant Molec Biol 56(2):225–239.

49 Lee K & Seo PJ (2017) Coordination of matrix attachment and ATP-dependent chromatin remodeling regulate auxin biosynthesis and Arabidopsis hypocotyl elongation. Plos One 12(7).

50 Yachie N, Saito R, Sugahara J, Tomita M, & Ishihama Y (2009) In Silico Analysis of Phosphoproteome Data Suggests a Rich-get-richer Process of Phosphosite Accumulation over Evolution. Molecular & Cellular Proteomics 8(5):1061–1071.

51 Favero DS, et al. (2016) SUPPRESSOR OF PHYTOCHROME B4-#3 Represses Genes Associated with Auxin Signaling to Modulate Hypocotyl Growth. Plant Physiol 171(4):2701–2716.

52 Xiao CW, Chen FL, Yu XH, Lin CT, & Fu YF (2009) Over-expression of an AT-hook gene, AHL22, delays flowering and inhibits the elongation of the hypocotyl in Arabidopsis thaliana. Plant Molec Biol 71(1-2):39–50.

53 Jiang H, et al. (2017) Ectopic application of the repressive histone modification H3K9me2 establishes post-zygotic reproductive isolation in Arabidopsis thaliana. Genes Develop 31(12):1272–1287.

54 Bigeard J, Rayapuram N, Bonhomme L, Hirt H, & Pflieger D (2014) Proteomic and phosphoproteomic analyses of chromatin-associated proteins from Arabidopsis thaliana. Proteomics 14(19):2141–2155.

55 Zhou LG, et al. (2016) A novel gene OsAHL1 improves both drought avoidance and drought tolerance in rice. Scientific Reports 6: 30264

56 Skirycz A, et al. (2011) Survival and growth of Arabidopsis plants given limited water are not equal. Nat Biotech 29(3):212–214.

57 Wang K, et al. (2018) Two Abscisic Acid Responsive Plastid Lipase Genes Involved in Jasmonic Acid Biosynthesis in Arabidopsis thaliana. Plant Cell 30(5): 1006–1022.

58 Bhaskara GB, Yang T-H, & Verslues PE (2015) Dynamic proline metabolism: Importance and regulation in water limited environments. Frontiers in Plant Science 6: 484.

59 Verslues PE, Agarwal M, Katiyar-Agarwal S, Zhu J, & Zhu JK (2006) Methods and concepts in quantifying resistance to drought, salt and freezing, abiotic stresses that affect plant water status. (vol 45, pg 523, 2006). Plant Journal 45 (4):523–539.

60 Tsuda K, et al. (2012) An efficient Agrobacterium-mediated transient transformation of Arabidopsis. Plant Journal 69(4):713–719.

61 Grefen C & Blatt MR (2012) A 2in1 cloning system enables ratiometric bimolecular fluorescence complementation (rBiFC). Biotechniques 53(5):311–314.

