## Supplementary material for "Phosphoproteomics of Arabidopsis Highly ABA-Induced1 identifies AT-Hook Like10 phosphorylation required for stress growth regulation"

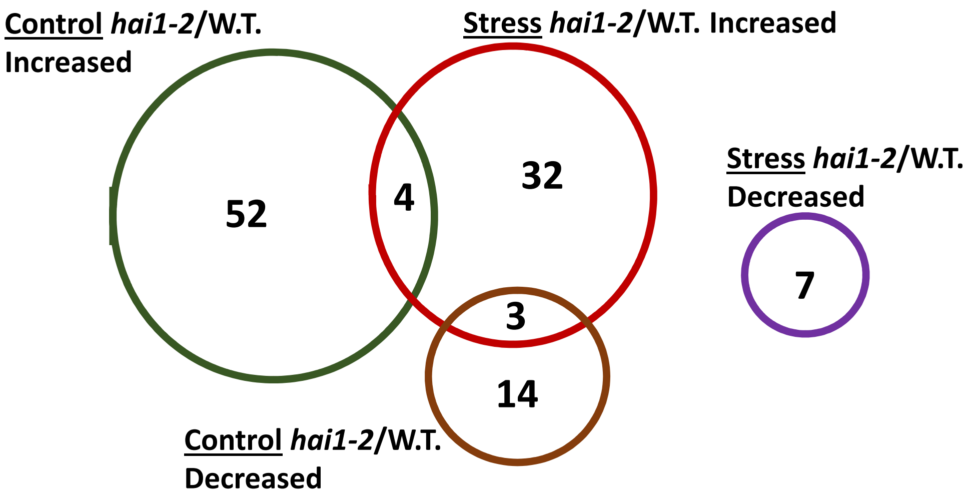


**4 Phosphopeptides increased in *hai1-2* control and stress:**

**AT5G64690.1** neurofilament triplet H protein-like protein

**AT1G35580.1** CYTOSOLIC INVERTASE 1 (CINV1);ALKALINE/NEUTRAL INVERTASE G (A/N-InvG), Interacts with PIP5K9.

**AT1G17210.1** IAP-LIKE PROTEIN 1 (ILP1)

**AT1G51140.1** FLOWERING BHLH 3 (FBH3);ABA-RESPONSIVE KINASE SUBSTRATE 1 (AKS1);ATCFL1 ASSOCIATED PROTEIN 1 (CFLAP1). Basic helix-loop-helix-type transcription factor, DNA-binding capacity is inhibited in response to ABA through phosphorylation-dependent monomerization.

**3 Phosphopeptides increased in *hai1-2* stress and decreased in *hai1-2* control:**

**AT3G55460.1** SC35-LIKE SPLICING FACTOR 30 (At-SCL30);SC35-LIKE SPLICING FACTOR 30 (SCL30). SC35-like splicing factor that is localized to nuclear speckles

**AT2G42670.2** PLANT CYSTEINE OXIDASE 4 (PCO4)

**AT4G15020.1** hAT transposon superfamily

**SI Appendix Figure S1: Overlap between proteins with phosphopeptides of altered abundance in *hai1-2* relative to wild type (W.T.) in the control and stress treatments.** Complete lists of phosphopeptides of significantly increased or decreased abundance in *hai1-2* relative to wild type in control and low water potential stress can be found in Datasets S2 and S3.

#### A *hai1-2*/W.T. Control

##### Increased abundance phosphopeptides

| # | Motif | Motif Score | Foreground Matches | Foreground Size | Background Matches | Background Size | Fold Increase |
|---|---|---|---|---|---|---|---|
| 1. | .....M.R..s.......... | 15.55 | 6 | 56 | 1357 | 1013205 | 80.00 |
| 2. | ..........sP......N.. | 12.75 | 5 | 50 | 2263 | 1011848 | 44.71 |
| 3. | .......R..sL......... | 8.58 | 5 | 45 | 5340 | 1009585 | 21.01 |
| 4. | ..........sP......... | 6.39 | 12 | 40 | 50733 | 1004245 | 5.94 |
| 5. | .......R..s.......... | 3.54 | 7 | 28 | 45887 | 953512 | 5.19 |
| 6. | ..........s...E...... | 2.84 | 6 | 21 | 56953 | 907625 | 4.55 |
| 7. | ..K.......s.......... | 2.78 | 5 | 15 | 53103 | 850672 | 5.34 |

1 2 3 4 5 6 7

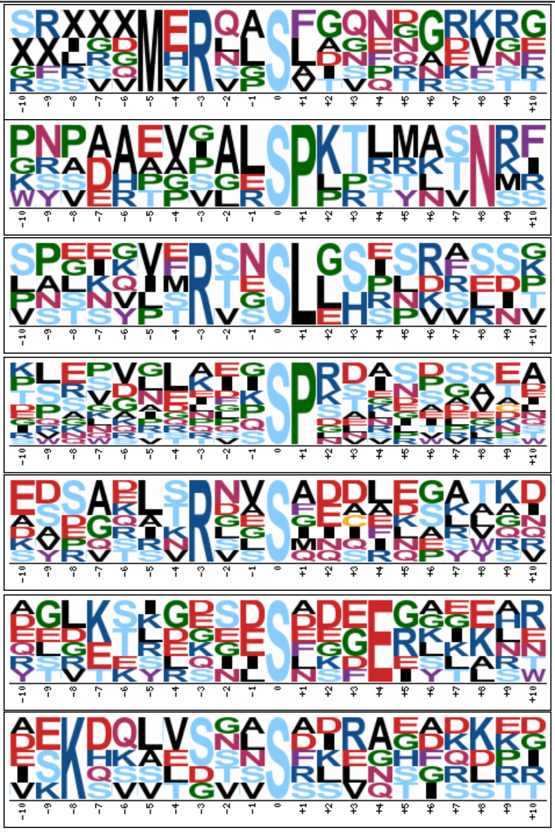

##### Decreased abundance phosphopeptides

| # | Motif | Motif Score | Foreground Matches | Foreground Size | Background Matches | Background Size | Fold Increase |
|---|---|---|---|---|---|---|---|
| 1. | ..........sP......... | 6.05 | 8 | 17 | 53062 | 1013205 | 8.99 |

1

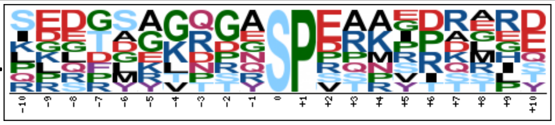

#### B *hai1-2*/W.T. Stress

##### Increased abundance phosphopeptides

| # | Motif | Motif Score | Foreground Matches | Foreground Size | Background Matches | Background Size | Fold Increase |
|---|---|---|---|---|---|---|---|
| 1. | .R........sP......... | 17.23 | 6 | 44 | 2839 | 1013205 | 48.67 |
| 2. | ..........sP......... | 8.76 | 14 | 38 | 50223 | 1010366 | 7.41 |
| 3. | .......R..s.......... | 5.57 | 9 | 24 | 52518 | 960143 | 6.86 |
| 4. | ..........s.......E.. | 2.72 | 5 | 15 | 58189 | 907625 | 5.20 |

1 2 3 4

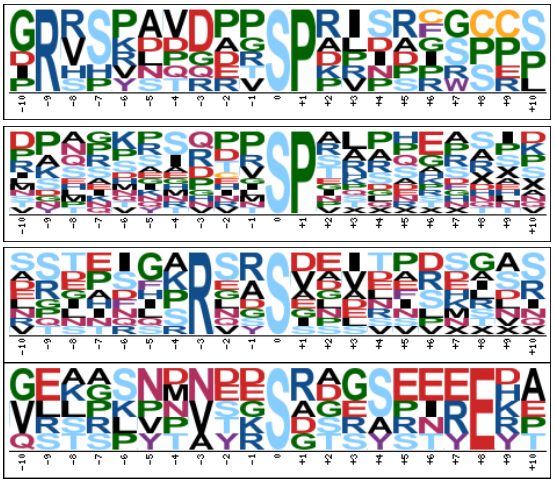

##### Decreased abundance phosphopeptides

| # | Motif | Motif Score | Foreground Matches | Foreground Size | Background Matches | Background Size | Fold Increase |
|---|---|---|---|---|---|---|---|
| 1. | ..........sP......... | 5.82 | 6 | 9 | 53062 | 1013205 | 12.73 |

1

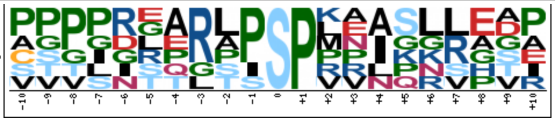

**SI Appendix Figure S2: Motif analysis of phosphopeptides of increased or decreased abundance in *hai1-2* relative to wild type.** Pre-aligned 21-amino acid sequences (Dataset S4) were submitted to Motif-X to discover overrepresented motifs (analysis parameters: minimum number of occurrences = 5, significance = 0.01, background = Arabidopsis proteome).

A. Motifs found in phosphopeptides of increased or decreased abundance in *hai1-2* in the unstressed control.

B. Motifs found in phosphopeptides of increased or decreased abundance in *hai1-2* under low water potential stress.

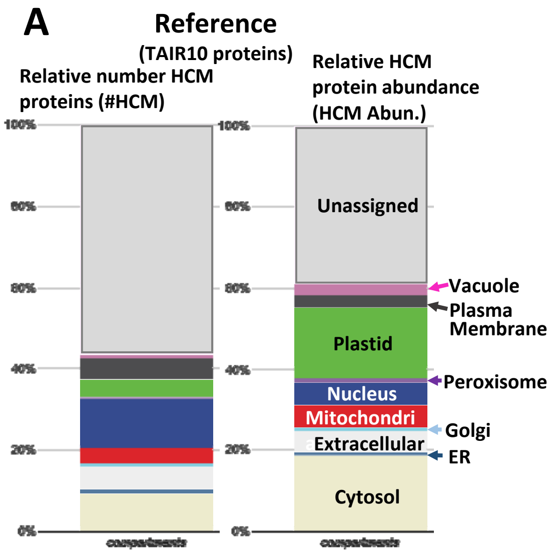


**SI Appendix Figure S3: Subcellular localization analysis of putative HAI1 target proteins shows an enrichment of nuclear proteins under stress**
Proteins with increased or decreased phosphopeptide abundance in *hai1-2* were submitted to the Multiple Marker Abundance Profiling (MMAP) tool in SUBA (http://suba.live/toolbox-app.html ). This tool searches for High Confidence Marker (HCM) proteins and analyzes both their number and relative protein abundance compared to other proteins in the submitted data.
**A.** Reference data showing localization and relative abundance of all proteins from TAIR10 annotation.
**B.** Profiles of HCM localization and abundance for proteins with increased or decreased phosphopeptide abundance in *hai1-2* in the unstressed control (Dataset S2).
**C.** Profiles of HCM localization and abundance for proteins with altered phosphopeptide abundance in *hai1-2* under stress (Dataset S3)

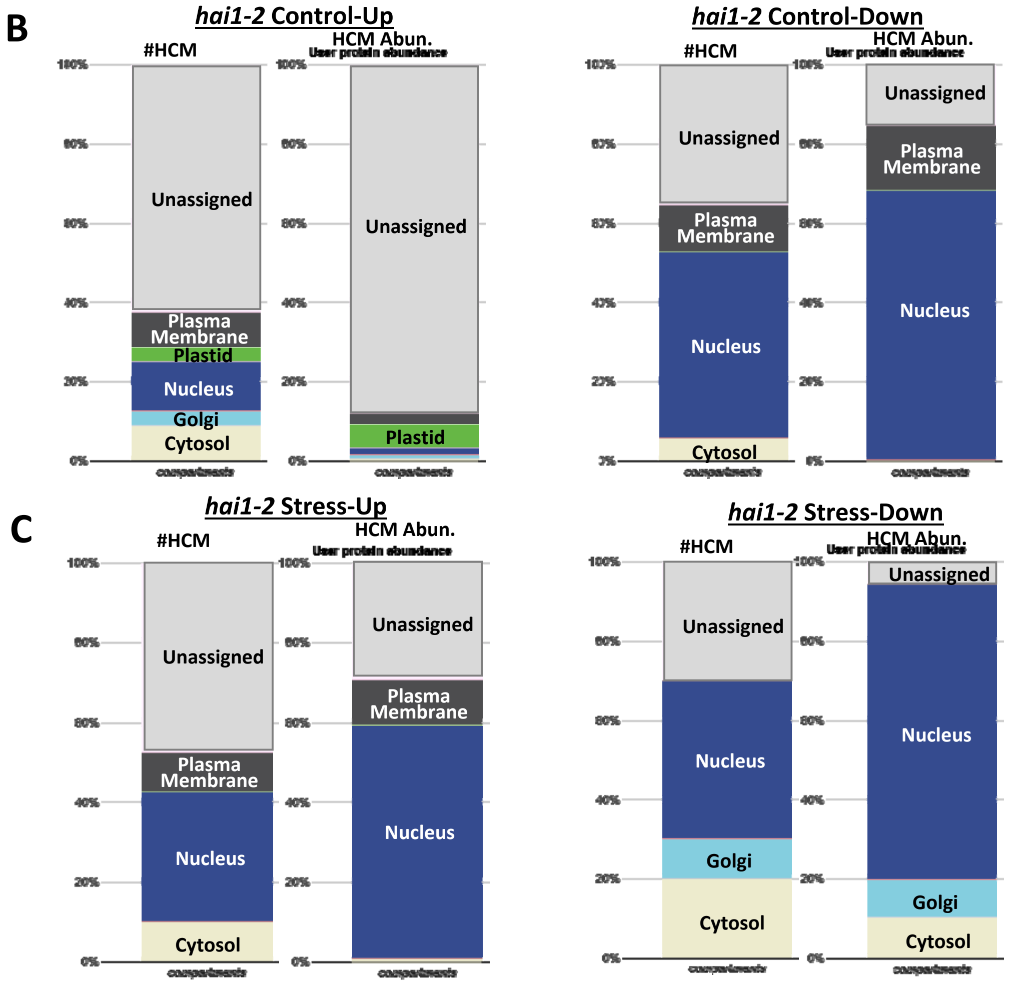

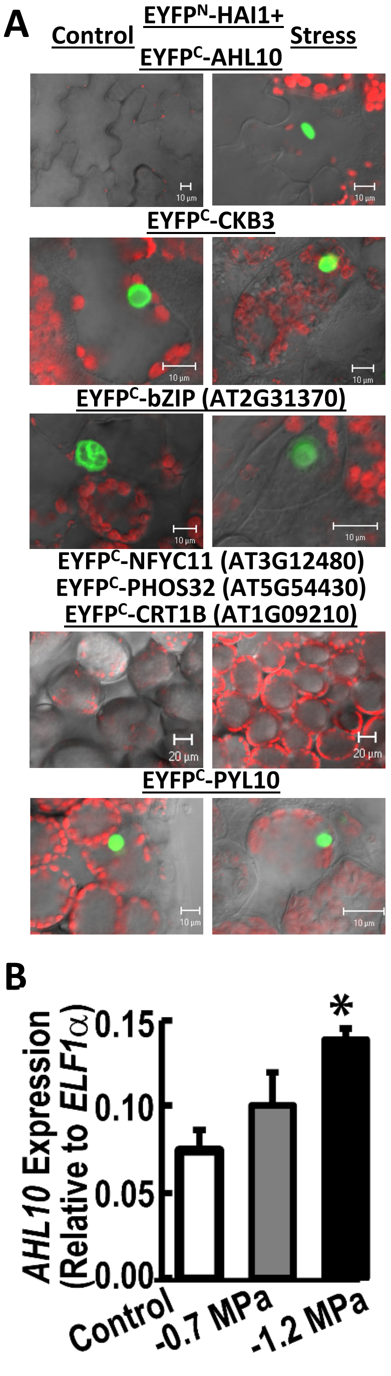


**SI Appendix Figure S4: BiFC protein interaction assays for HAI1 and candidate proteins identified by phosphoproteomics.**

**A.** HAI1 and the protein of interest were cloned into the pSITE-nEYFP-C1 and pSITE-cEYFP-C1 vectors, respectively. The constructs were transiently expressed in Arabidopsis seedlings which were then kept under unstressed conditions or exposed to low water potential stress for 72 h. Seven candidate proteins selected based on having increased phosphopeptide abundance in *hai1-2* under either control or stress conditions were tested. Note that for some of these candidates (CRT1B, NFYC11, bZIP) the increase in phosphopeptide abundance was relatively large but not significant in the statistical analysis.

Images shown are leaf mesophyll cells. Green color indicates the BiFC signal while red is chlorophyll autoflourescence. Scale bars show 10 or 20 $\mu$m as indicated in each panel. Only one representative image is shown for candidates where interaction could not be observed.

BiFC interaction was seen for HAI1 with AHL10, CKB3 and a bZIP protein. The HAI1-AHL10 interaction was only observed under stress. The interactions were observed in the nucleus consistent with the expected localization of each protein. No interaction was seen for CRT1B, PHOS32 or NFYC11. The known interaction of HAI1 with PYL10 was assayed as a positive control.

B. *AHL10* gene expression in wild type for unstressed control or 96 h after transfer to two different severities of low $y_w$ stress (-0.7 and -1.2 MPa). Data are means ± SE, n = 3. Asterisk (*) indicates significant difference compared to control (P ≤0.05 by T-test).

### SI Appendix Figure S5: Spectra of AHL10 phosphopeptides.

#### A. AHL10 (AT2G33620) S313 phosphorylation

**Sequence: vAPTQVLMTPsSPQSR,**

V1-iTRAQ8plex (304.20536 Da), S11-Phospho (79.96633 Da), Charge: +3, Monoisotopic m/z: 695.02277 Da (-0.79 mmu/-1.13 ppm), MH+: 2083.05375 Da, RT: 50.85 min, Identified with: SEQUEST (v1.20); XCorr: 5.16, Probability (FDR q-value): 0.00, Ions matched by search engine: 23/88, Fragment match tolerance used for search: 50 mmu

| #1 | b⁺ | b²⁺ | b⁺-P | Seq. | y⁺ | y²⁺ | y⁺-P | #2 |
|---|---|---|---|---|---|---|---|---|
| 1 | 404.28106 | 202.64417 | | V-iTRAQ8plex | | | | 16 |
| 2 | 475.31818 | 238.16273 | | A | 1679.78232 | 840.39480 | 1581.80543 | 15 |
| 3 | 572.37095 | 286.68911 | | P | 1608.74520 | 804.87624 | 1510.76831 | 14 |
| 4 | 673.41863 | 337.21295 | | T | 1511.69243 | 756.34985 | 1413.71554 | 13 |
| 5 | 801.47721 | 401.24224 | | Q | 1410.64475 | 705.82601 | 1312.66786 | 12 |
| 6 | 900.54563 | 450.77645 | | V | 1282.58617 | 641.79672 | 1184.60928 | 11 |
| 7 | 1013.62970 | 507.31849 | | L | 1183.51775 | 592.26251 | 1085.54086 | 10 |
| 8 | 1144.67020 | 572.83874 | | M | 1070.43368 | 535.72048 | 972.45679 | 9 |
| 9 | 1245.71788 | 623.36258 | | T | 939.39318 | 470.20023 | 841.41629 | 8 |
| 10 | 1342.77065 | 671.88896 | | P | 838.34550 | 419.67639 | 740.36861 | 7 |
| 11 | 1509.76901 | 755.38814 | 1411.79211 | S-Phospho | 741.29273 | 371.15000 | 643.31584 | 6 |
| 12 | 1596.80104 | 798.90416 | 1498.82414 | S | 574.29437 | 287.65082 | | 5 |
| 13 | 1693.85381 | 847.43054 | 1595.87691 | P | 487.26234 | 244.13481 | | 4 |
| 14 | 1821.91239 | 911.45983 | 1723.93549 | Q | 390.20957 | 195.60842 | | 3 |
| 15 | 1908.94442 | 954.97585 | 1810.96752 | S | 262.15099 | 131.57913 | | 2 |
| 16 | | | | R | 175.11896 | 88.06312 | | 1 |

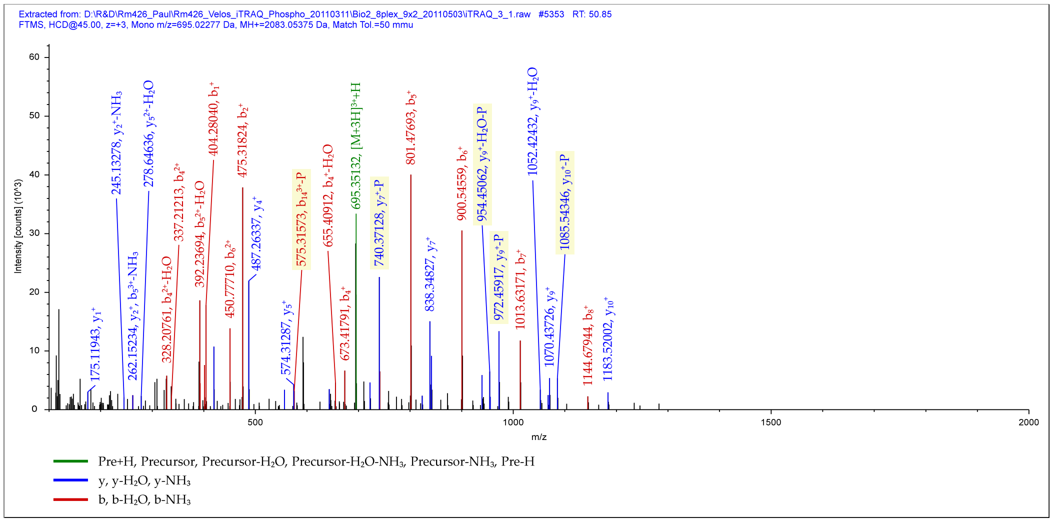


#### B. AHL10 (AT2G33620) S314 phosphorylation

**Sequence: vAPTQVLmTPSsPQSR,**

V1-iTRAQ8plex (304.20536 Da), M8-Oxidation (15.99492 Da), S12-Phospho (79.96633 Da), Charge: +3, Monoisotopic m/z: 700.35394 Da (-1.25 mmu/-1.78 ppm), MH+: 2099.04728 Da, RT: 42.91 min, Identified with: SEQUEST (v1.20); XCorr:5.54, Probability (FDR q-value): 0.00, Ions matched by search engine: 26/88, Fragment match tolerance used for search: 50 mmu

| #1 | b⁺ | b²⁺ | b⁺-P | Seq. | y⁺ | y²⁺ | y⁺-P | #2 |
|---|---|---|---|---|---|---|---|---|
| 1 | 404.28106 | 202.64417 | | V-iTRAQ8plex | | | | 16 |
| 2 | 475.31818 | 238.16273 | | A | 1695.77724 | 848.39226 | 1597.80034 | 15 |
| 3 | 572.37095 | 286.68911 | | P | 1624.74012 | 812.87370 | 1526.76322 | 14 |
| 4 | 673.41863 | 337.21295 | | T | 1527.68735 | 764.34731 | 1429.71045 | 13 |
| 5 | 801.47721 | 401.24224 | | Q | 1426.63967 | 713.82347 | 1328.66277 | 12 |
| 6 | 900.54563 | 450.77645 | | V | 1298.58109 | 649.79418 | 1200.60419 | 11 |
| 7 | 1013.62970 | 507.31849 | | L | 1199.51267 | 600.25997 | 1101.53577 | 10 |
| 8 | 1160.66511 | 580.83619 | | M-Oxidation | 1086.42860 | 543.71794 | 988.45170 | 9 |
| 9 | 1261.71279 | 631.36003 | | T | 939.39318 | 470.20023 | 841.41629 | 8 |
| 10 | 1358.76556 | 679.88642 | | P | 838.34550 | 419.67639 | 740.36861 | 7 |
| 11 | 1445.79759 | 723.40243 | | S | 741.29273 | 371.15000 | 643.31584 | 6 |
| 12 | 1612.79595 | 806.90161 | 1514.81906 | S-Phospho | 654.26070 | 327.63399 | 556.28381 | 5 |
| 13 | 1709.84872 | 855.42800 | 1611.87183 | P | 487.26234 | 244.13481 | | 4 |
| 14 | 1837.90730 | 919.45729 | 1739.93041 | Q | 390.20957 | 195.60842 | | 3 |
| 15 | 1924.93933 | 962.97330 | 1826.96244 | S | 262.15099 | 131.57913 | | 2 |
| 16 | | | | R | 175.11896 | 88.06312 | | 1 |

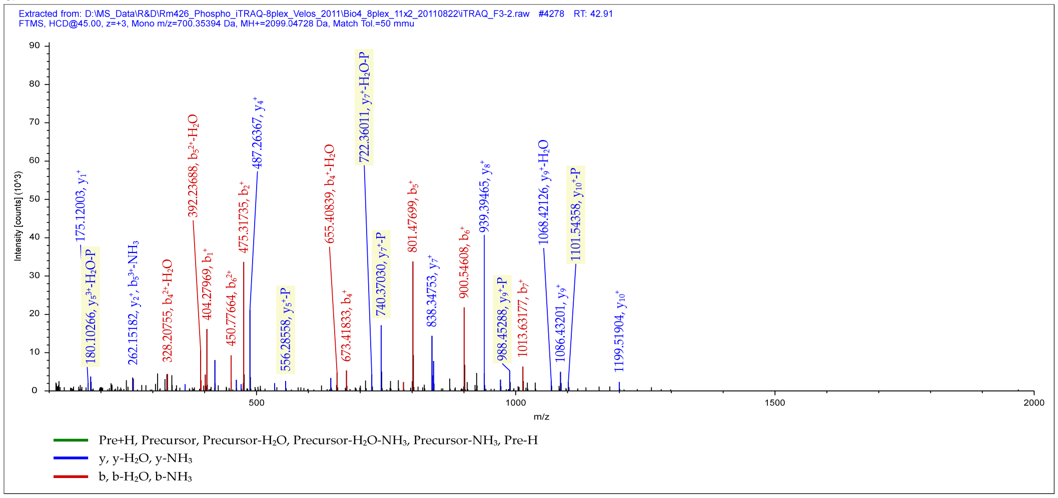

#### SI Appendix Figure S6: AHL13 phosphopeptide spectra.

**A. Sequence: aQNTPEPASAPANmLSFGGVGGPGsPR**, A1-iTRAQ8plex (304.20536 Da), M14-Oxidation (15.99492 Da), S25-Phospho (79.96633 Da), **Charge: +4**, Monoisotopic m/z: 742.60388 Da (-0.94 mmu/-1.27 ppm), MH+: 2967.39370 Da, RT: 49.06 min, Identified with: SEQUEST (v1.20); XCorr:8.23, Probability (FDR q-value): 0.00, Ions matched by search engine: 38/231, Fragment match tolerance used for search: 50 mmu

| #1 | $b^+$ | $b^{2+}$ | $b^+$-P | $b^{2+}$-P | Seq. | $y^+$ | $y^{2+}$ | $y^+$-P | $y^{2+}$-P | #2 |
|---|---|---|---|---|---|---|---|---|---|---|
| 1 | 376.24976 | 188.62852 | | | A-iTRAQ8plex | | | | | 27 |
| 2 | 504.30834 | 252.65781 | | | Q | 2592.15499 | 1296.58113 | 2494.17809 | 1247.59268 | 26 |
| 3 | 618.35127 | 309.67927 | | | N | 2464.09641 | 1232.55184 | 2366.11951 | 1183.56339 | 25 |
| 4 | 719.39895 | 360.20311 | | | T | 2350.05348 | 1175.53038 | 2252.07658 | 1126.54193 | 24 |
| 5 | 816.45172 | 408.72950 | | | P | 2249.00580 | 1125.00654 | 2151.02890 | 1076.01809 | 23 |
| 6 | 945.49432 | 473.25080 | | | E | 2151.95303 | 1076.48015 | 2053.97613 | 1027.49170 | 22 |
| 7 | 1042.54709 | 521.77718 | | | P | 2022.91043 | 1011.95885 | 1924.93353 | 962.97040 | 21 |
| 8 | 1113.58421 | 557.29574 | | | A | 1925.85766 | 963.43247 | 1827.88076 | 914.44402 | 20 |
| 9 | 1200.61624 | 600.81176 | | | S | 1854.82054 | 927.91391 | 1756.84364 | 878.92546 | 19 |
| 10 | 1271.65336 | 636.33032 | | | A | 1767.78851 | 884.39789 | 1669.81161 | 835.40944 | 18 |
| 11 | 1368.70613 | 684.85670 | | | P | 1696.75139 | 848.87933 | 1598.77449 | 799.89088 | 17 |
| 12 | 1439.74325 | 720.37526 | | | A | 1599.69862 | 800.35295 | 1501.72172 | 751.36450 | 16 |
| 13 | 1553.78618 | 777.39673 | | | N | 1528.66150 | 764.83439 | 1430.68460 | 715.84594 | 15 |
| 14 | 1700.82159 | 850.91443 | | | M-Oxidation | 1414.61857 | 707.81292 | 1316.64167 | 658.82447 | 14 |
| 15 | 1813.90566 | 907.45647 | | | L | 1267.58315 | 634.29521 | 1169.60626 | 585.30677 | 13 |
| 16 | 1900.93769 | 950.97248 | | | S | 1154.49908 | 577.75318 | 1056.52219 | 528.76473 | 12 |
| 17 | 2048.00611 | 1024.50669 | | | F | 1067.46705 | 534.23716 | 969.49016 | 485.24872 | 11 |
| 18 | 2105.02758 | 1053.01743 | | | G | 920.39863 | 460.70295 | 822.42174 | 411.71451 | 10 |
| 19 | 2162.04905 | 1081.52816 | | | G | 863.37716 | 432.19222 | 765.40027 | 383.20377 | 9 |
| 20 | 2261.11747 | 1131.06237 | | | V | 806.35569 | 403.68148 | 708.37880 | 354.69304 | 8 |
| 21 | 2318.13894 | 1159.57311 | | | G | 707.28727 | 354.14727 | 609.31038 | 305.15883 | 7 |
| 22 | 2375.16041 | 1188.08384 | | | G | 650.26580 | 325.63654 | 552.28891 | 276.64809 | 6 |
| 23 | 2472.21318 | 1236.61023 | | | P | 593.24433 | 297.12580 | 495.26744 | 248.13736 | 5 |
| 24 | 2529.23465 | 1265.12096 | | | G | 496.19156 | 248.59942 | 398.21467 | 199.61097 | 4 |
| 25 | 2696.23301 | 1348.62014 | 2598.25612 | 1299.63170 | S-Phospho | 439.17009 | 220.08868 | 341.19320 | 171.10024 | 3 |
| 26 | 2793.28578 | 1397.14653 | 2695.30889 | 1348.15808 | P | 272.17173 | 136.58950 | | | 2 |
| 27 | | | | | R | 175.11896 | 88.06312 | | | 1 |

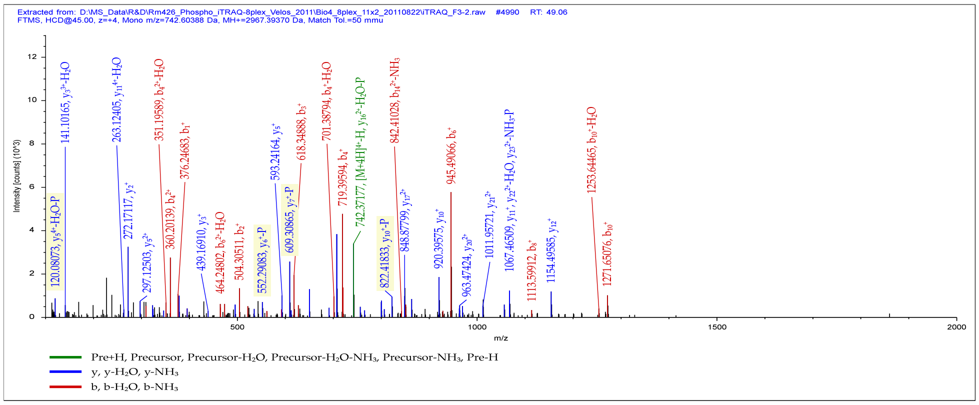


**B. Sequence: aQNTPEPASAPANmLSFGGVGGPGsPR**, A1-iTRAQ8plex (304.20536 Da), M14-Oxidation (15.99492 Da), S25-Phospho (79.96633 Da), **Charge: +3**, Monoisotopic m/z: 989.80469 Da (+0.68 mmu/+0.69 ppm), MH+: 2967.39951 Da, RT: 48.72 min, Identified with: SEQUEST (v1.20); XCorr:7.00, Probability (FDR q-value): 0.00, Ions matched by search engine: 35/154, Fragment match tolerance used for search: 50 mmu

| #1 | $b^+$ | $b^{2+}$ | $b^+$-P | $b^{2+}$-P | Seq. | $y^+$ | $y^{2+}$ | $y^+$-P | $y^{2+}$-P | #2 |
|---|---|---|---|---|---|---|---|---|---|---|
| 1 | 376.24976 | 188.62852 | | | A-iTRAQ8plex | | | | | 27 |
| 2 | 504.30834 | 252.65781 | | | Q | 2592.15499 | 1296.58113 | 2494.17809 | 1247.59268 | 26 |
| 3 | 618.35127 | 309.67927 | | | N | 2464.09641 | 1232.55184 | 2366.11951 | 1183.56339 | 25 |
| 4 | 719.39895 | 360.20311 | | | T | 2350.05348 | 1175.53038 | 2252.07658 | 1126.54193 | 24 |
| 5 | 816.45172 | 408.72950 | | | P | 2249.00580 | 1125.00654 | 2151.02890 | 1076.01809 | 23 |
| 6 | 945.49432 | 473.25080 | | | E | 2151.95303 | 1076.48015 | 2053.97613 | 1027.49170 | 22 |
| 7 | 1042.54709 | 521.77718 | | | P | 2022.91043 | 1011.95885 | 1924.93353 | 962.97040 | 21 |
| 8 | 1113.58421 | 557.29574 | | | A | 1925.85766 | 963.43247 | 1827.88076 | 914.44402 | 20 |
| 9 | 1200.61624 | 600.81176 | | | S | 1854.82054 | 927.91391 | 1756.84364 | 878.92546 | 19 |
| 10 | 1271.65336 | 636.33032 | | | A | 1767.78851 | 884.39789 | 1669.81161 | 835.40944 | 18 |
| 11 | 1368.70613 | 684.85670 | | | P | 1696.75139 | 848.87933 | 1598.77449 | 799.89088 | 17 |
| 12 | 1439.74325 | 720.37526 | | | A | 1599.69862 | 800.35295 | 1501.72172 | 751.36450 | 16 |
| 13 | 1553.78618 | 777.39673 | | | N | 1528.66150 | 764.83439 | 1430.68460 | 715.84594 | 15 |
| 14 | 1700.82159 | 850.91443 | | | M-Oxidation | 1414.61857 | 707.81292 | 1316.64167 | 658.82447 | 14 |
| 15 | 1813.90566 | 907.45647 | | | L | 1267.58315 | 634.29521 | 1169.60626 | 585.30677 | 13 |
| 16 | 1900.93769 | 950.97248 | | | S | 1154.49908 | 577.75318 | 1056.52219 | 528.76473 | 12 |
| 17 | 2048.00611 | 1024.50669 | | | F | 1067.46705 | 534.23716 | 969.49016 | 485.24872 | 11 |
| 18 | 2105.02758 | 1053.01743 | | | G | 920.39863 | 460.70295 | 822.42174 | 411.71451 | 10 |
| 19 | 2162.04905 | 1081.52816 | | | G | 863.37716 | 432.19222 | 765.40027 | 383.20377 | 9 |
| 20 | 2261.11747 | 1131.06237 | | | V | 806.35569 | 403.68148 | 708.37880 | 354.69304 | 8 |
| 21 | 2318.13894 | 1159.57311 | | | G | 707.28727 | 354.14727 | 609.31038 | 305.15883 | 7 |
| 22 | 2375.16041 | 1188.08384 | | | G | 650.26580 | 325.63654 | 552.28891 | 276.64809 | 6 |
| 23 | 2472.21318 | 1236.61023 | | | P | 593.24433 | 297.12580 | 495.26744 | 248.13736 | 5 |
| 24 | 2529.23465 | 1265.12096 | | | G | 496.19156 | 248.59942 | 398.21467 | 199.61097 | 4 |
| 25 | 2696.23301 | 1348.62014 | 2598.25612 | 1299.63170 | S-Phospho | 439.17009 | 220.08868 | 341.19320 | 171.10024 | 3 |
| 26 | 2793.28578 | 1397.14653 | 2695.30889 | 1348.15808 | P | 272.17173 | 136.58950 | | | 2 |
| 27 | | | | | R | 175.11896 | 88.06312 | | | 1 |

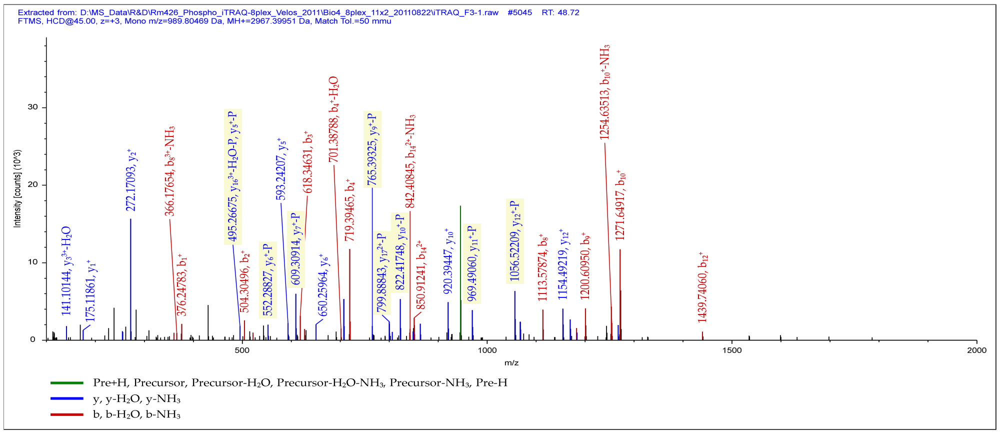

A

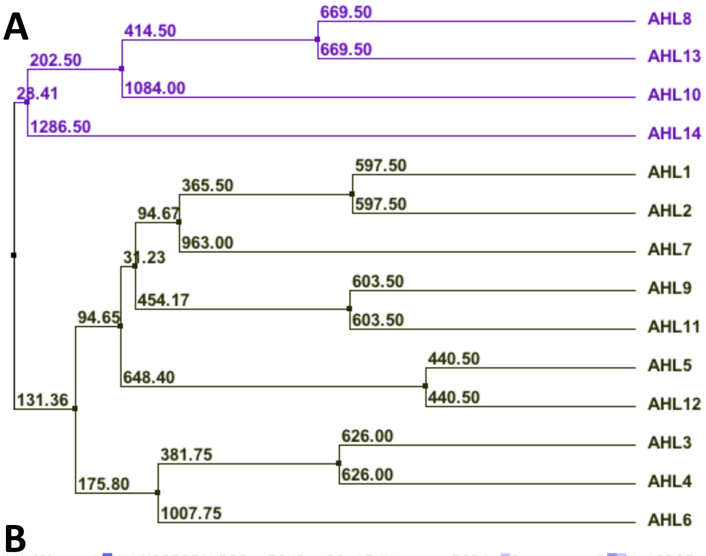


**SI Appendix Figure S7: Alignment of Clade B AHLs and location of AHL10/13 phosphorylation site.**

A. Phylogenetic tree of Clade B AHLs
B. Alignment of Clade B AHL protein sequences. The red box marks the AHL10 S313/S314 phosphorylation sites. Positions of the two AT-hook domains and PPC/DUF296 domain are also marked.

B

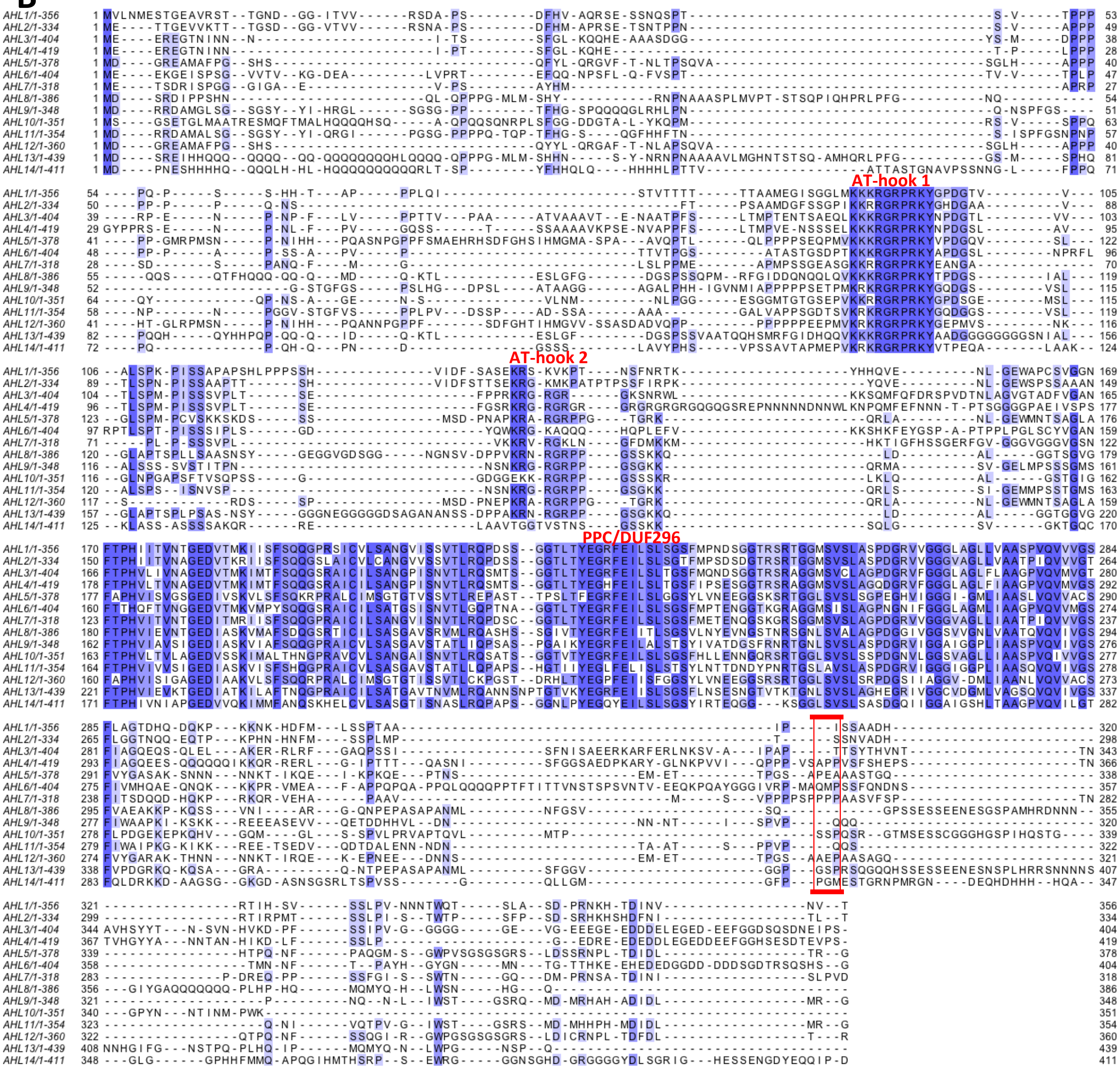

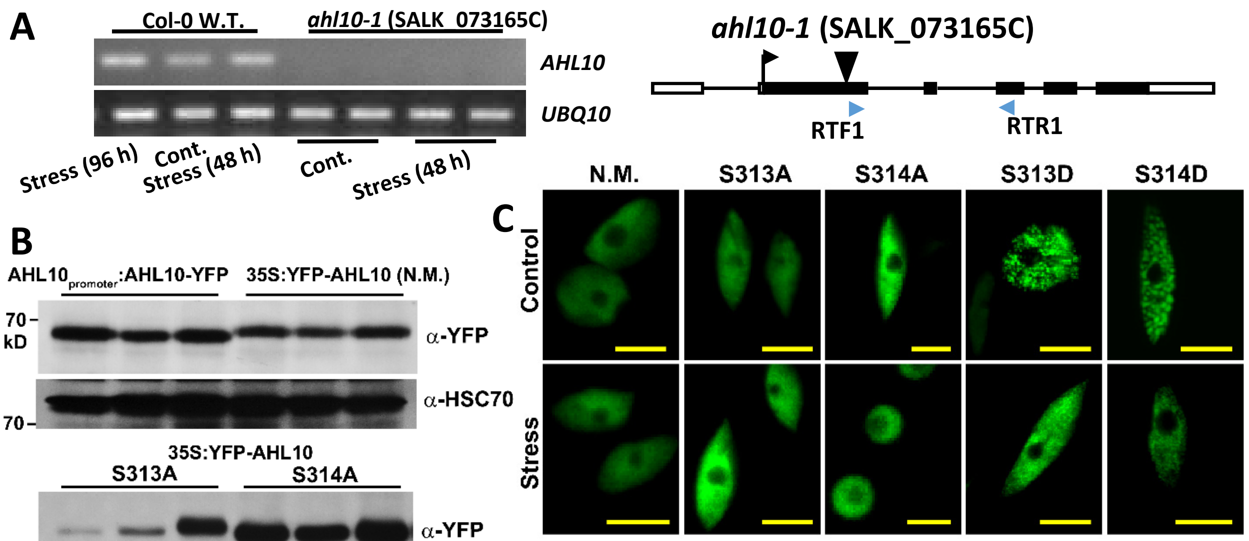


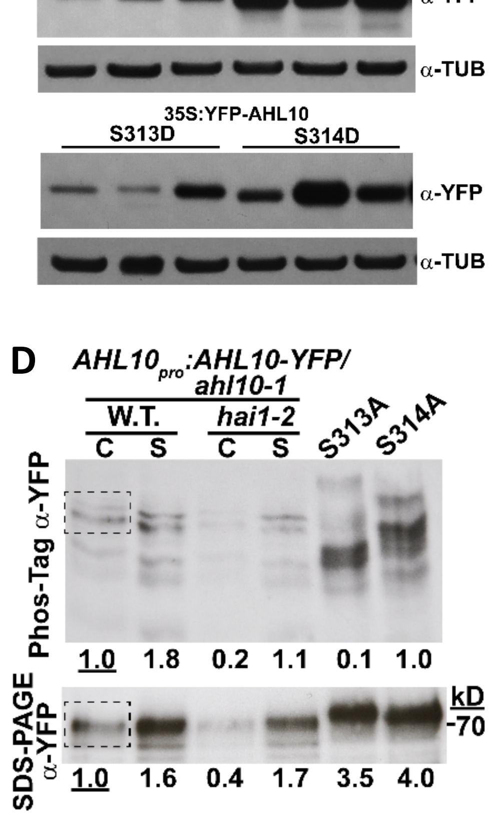


**SI Appendix Figure S8: *ahl10-1* characterization, AHL10 phosphonull and phosphomimic protein expression, localization and phosphorylation status.**

**A.** RT-PCR analysis of *ahl10-1* confirmed that it is a knockout mutant. Stress = seedlings transferred to --1.2 MPa. UBQ10 was amplified as a control for the RT-PCR. Diagram shows AHL10 gene structure, position of the T-DNA insertion in *ahl10-1* and position of the primers used for RT-PCR (black boxes indicate exons and unfilled boxes indicate UTR regions).

**B.** Immunoblot analysis to confirm protein expression in *AHL10* transgenic lines. Blots were stripped and reprobed with antisera recognizing HSC70 or Tubulin as a loading control. All samples were from unstressed seedlings. Note that all transgenic lines were constructed in the ahl10-1 mutant background and $AHL10_{promoter}$*:AHL10-YFP/ahl10* was also crossed into *hai1-2*.

**C.** Protein localization of AHL10 wild type (N.M., no mutation) as well as phosphonull (S313A, S314A) and phosphomimic (S313D, S314D) mutants. In all cases, AHL10 was seen exclusively in the nucleus. The phosphomimic versions of AHL10 partially localized in foci that were not seen in wild type or phosphonull AHL10. For all images, YFP-AHL10 was expressed under control of the 35S promoter in the *ahl10-1* background (*35S:YFP-AHL10/ahl10-1*) .

**D.** Immunoprecipitation of AHL10 from $AHL10_{promoter}$*:AHL10-YFP/ahl10-1 (*in W.T. or *hai1-2* background*)* plants from the unstressed control or -0.7 MPa stress treatment. 100 µg of total protein was used for each immunoprecipiation. For comparsion, AHL10 S3134 and S314A were immunoprecipitated from seedlings exposed to -0.7 MPa. The experiment was repeated with consistent results. The dashed box in the wild type control lane indicates the region selected from each lane for quantification of band intensities. Relative band intensities are indicated by the numbers below each lane.

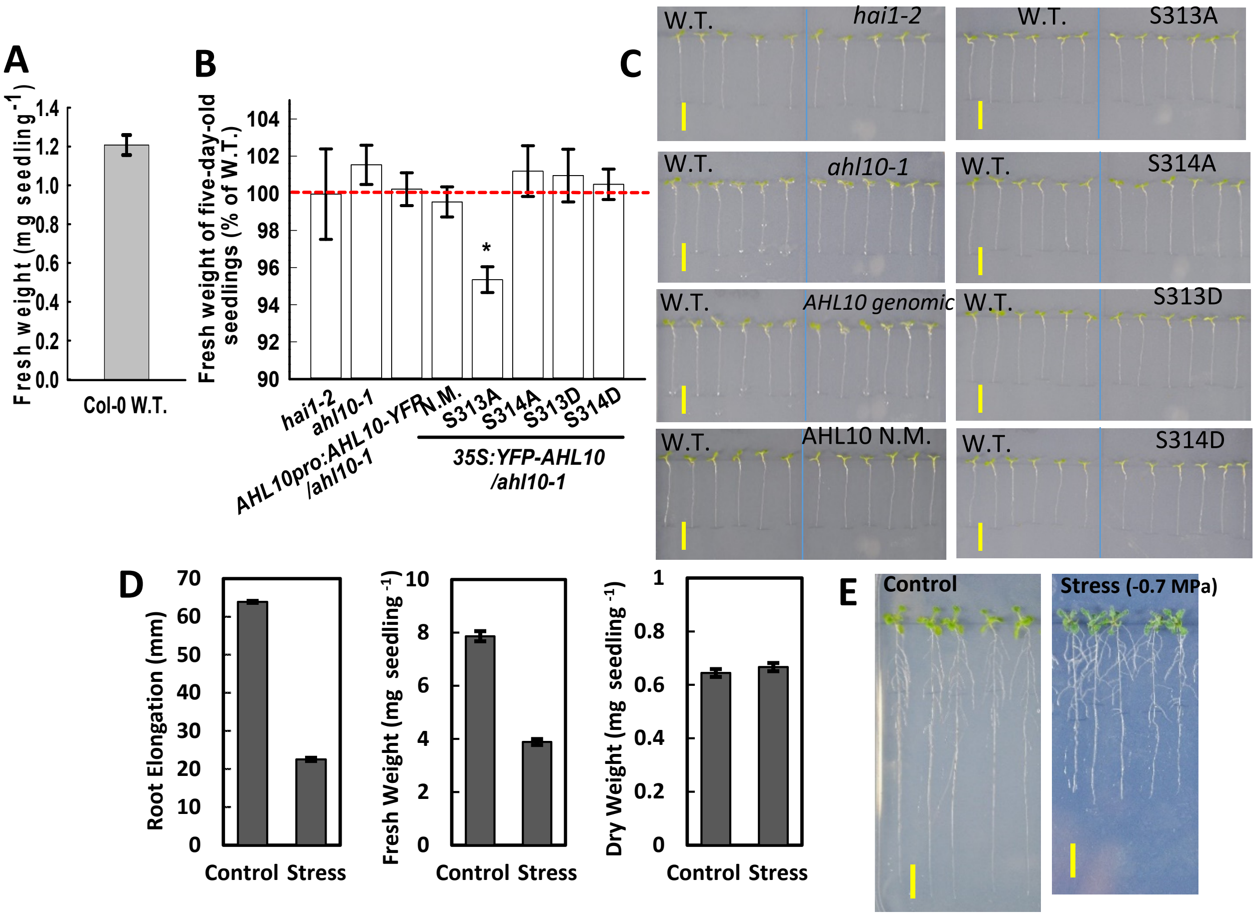


**SI Appendix Figure S9: Wild type growth data used to normalize the PEG-agar plate assay data for mutant and transgenic lines shown in Figure 2, fresh weight of 5-day-old seedlings before transfer to stress.**

A. Fresh weight of 5-day-old Col-0 W.T. seedlings at the time of transfer to stress or control treatments for PEG-plate growth assays. (mean ± SE, n = 280)

B. Fresh weights of five-day-old seedlings at the time of transfer to stress and control treatments in the PEG-plate growth assays. Data are relative to the Col-0 wild type (W.T.) grown on the same plates as the genotype assayed (mean ± SE, n = 35 to 40, combined from three independent experiments). Asterisk (*) indicates significant difference compared with the wild type by one-sided T-test ($P \le 0.05$). Dashed red line indicates the wild type level (100%).

C. Representative five-day-old seedlings on a half-strength MS agar plates. “AHL10 genomic” is *AHL10*$_{promoter}$*:AHL10-YFP/ahl10-1.* Other transgenic lines are non-mutated (N.M.), phosphonull (S313A, S314A) and phosphomimic (S313D, S314D) AHL10 expressed under control of 35S promoter in the *ahl10-1* mutant background (*35S:YFP-AHL10/ahl10-1*). Scale bars indicate 1cm.

D. Wild type (Col-0) root elongation, seedling fresh weight and dry weight. Five-day-old seedlings were transferred to either fresh control media (-0.25 MPa) or moderate severity low water potential stress (-0.7 MPa). Root elongation was measured over the following 5 days for control and 10 days for stress. Fresh weight and dry weigh were measured for groups of 5-6 seedlings collected at the end of the experiment. Data are combined from six representative experiments (mean ± S.E.; n = 435-650 for root elongation and 125-190 for fresh and dry weight). Comparison of this data to the fresh weight at the time of transfer (A) shows that the seedlings continued to grow and increase in fresh weight at -0.7 MPa but at a reduced rate compared to control. Seedlings allowed to grow for 10 days after transfer to -0.7 MPa were able to accumulate a similar total dry weight as unstressed seedlings harvested at 5 days after transfer.

E. Representative pictures of wild type seedlings under control or stress conditions. Photos were taken at the end of experiment (5 days after transfer for control, 10 days after transfer for stress). Black lines along the root indicate the position of the primary root apex at the time of transfer. Scale bars indicate 1 cm.

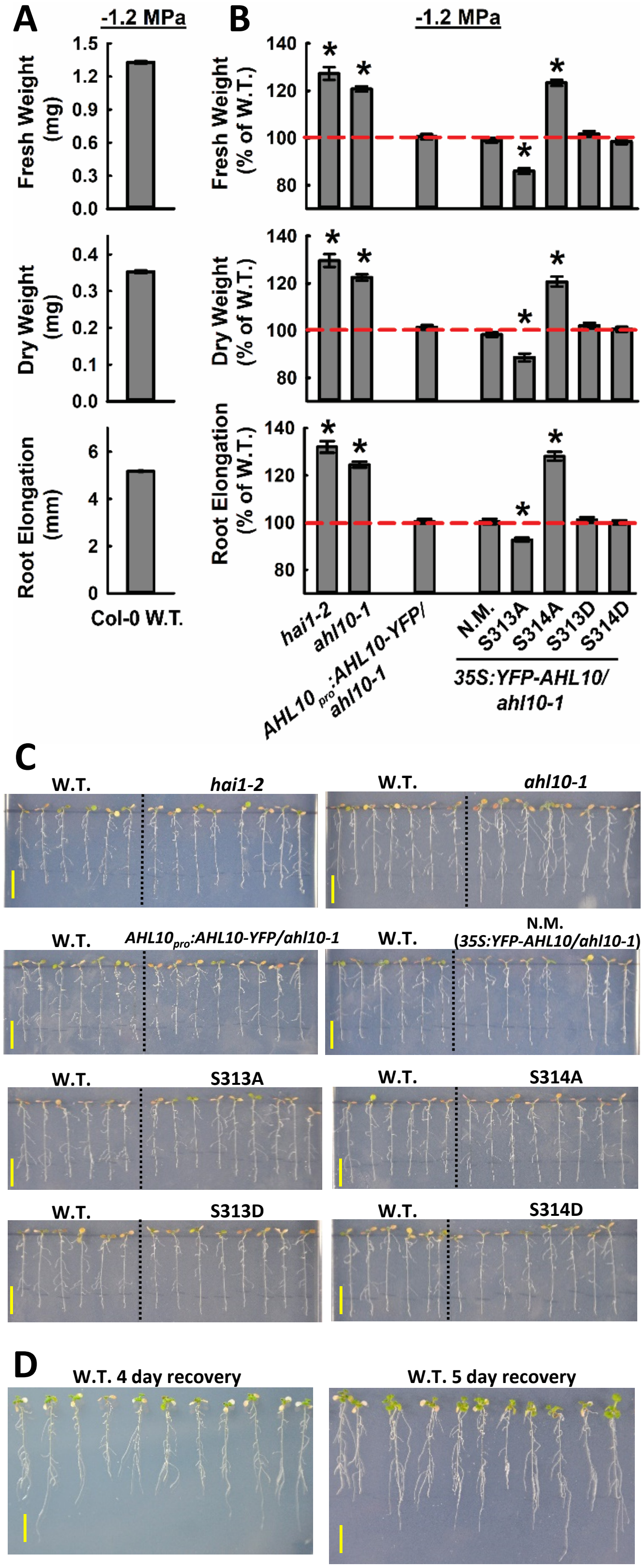


**SI Appendix Figure S10: Growth of AHL10 mutant and transgenic lines after transfer to -1.2 MPa PEG-agar plates.**

**A.** Fresh weight, dry weight and root elongation data of Col-0 W.T. used to normalize the data in B. Five-day-old seedlings were transferred to -1.2 MPa agar plates and growth parameters measured 10 days later. Data are mean ± SE, n = 139 for fresh weight and dry weight, n = 835 for root elongation, combined from two independent experiments.

**B.** Relative fresh weight, dry weight and root elongation of AHL10 and HAI1 mutant and transgenic lines 10 days after transfer of 5-day-old seedlings to -1.2 MPa agar plates. Data are presented relative to wild type seedlings assayed in the same PEG-agar plates (dashed red line indicates the wild type level, 100%). Data are mean ± SE, n = 6-10 combined from two independent experiments. Asterisk (*) indicates significant difference compared with the wild type by one-sample T-test ($P \leq 0.05$). Dashed red line indicates the wild type level (100%). N.M. = Non mutated (wild type) AHL10.

**C.** Representative images of Col-0 wild type (W.T.) and mutant/transgenic seedlings. Five day-old seedlings were transferred to -1.2 MPa agar plates for 10 days before photographs were taken. Images are from the same experiments shown in A and B. Scale bars indicate 1 cm. Black lines behind the roots indicated the position of the root apices at the time of transfer.

**D.** Representative images of seedlings subjected to -1.2 MPa stress for 10 days as described above and then transferred back to unstressed control media to determine the ability of the seedlings to recover and resume growth. Several independent experiments were performed and 100% of seedlings were alive and recovered in all cases. Photographs were taken 4 or 5 days after transfer back to the control media, as indicated. Scale bars indicate 1 cm.

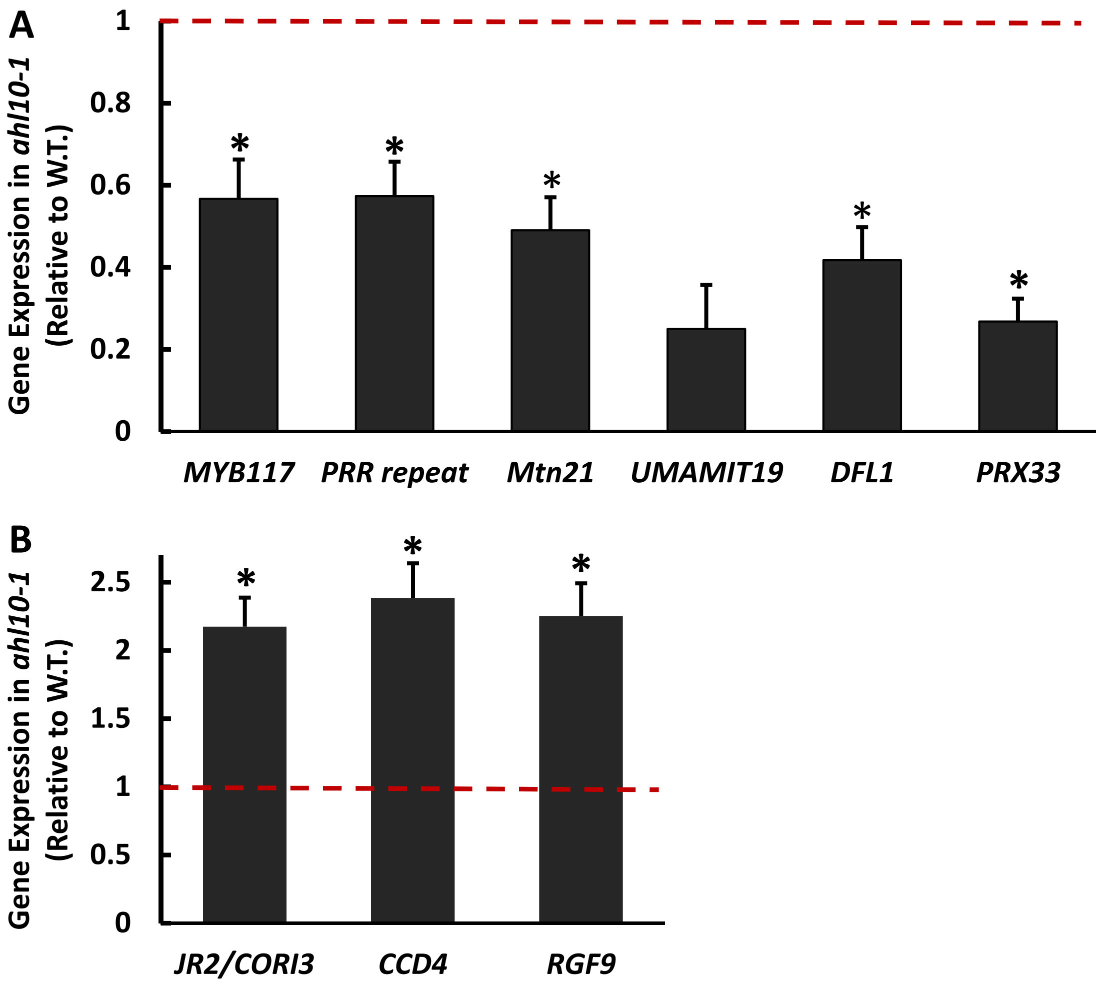


**SI Appendix Figure S11: Validation of additional gene expression changes in *ahl10-1* at low water potential.**

A. QPCR assay of genes found to be down-regulated in *ahl10-1* during stress by RNAseq (Dataset S9). Seedlings were transferred to -0.7 MPa for 96 h and gene expression assayed. Data are means ± S.E., n =3 from 3 independent experiments. Expression data are relative to wild type assayed in the same experiment. Red dashed line indicates the wild type expression level (set to 1). Asterisks (*) indicate significant difference relative to wild type (by one-sided T-test, P ≤ 0.05).

B. QPCR assay of genes found to be up-regulated in *ahl10-1* during stress by RNAseq (Dataset S9). Data format is the same as in A.

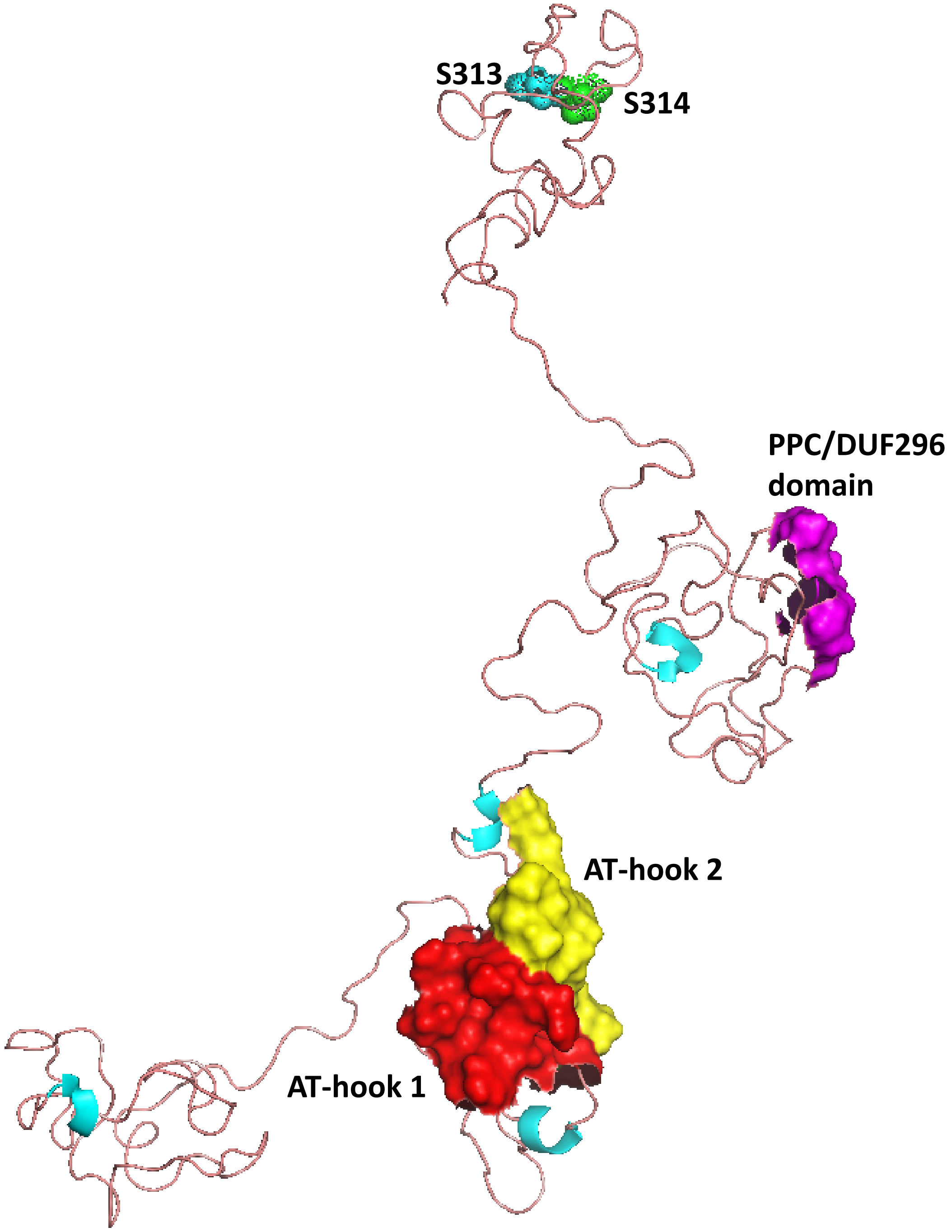


**SI Appendix Figure S12: Predicted AHL10 structure and location of the S313 and S314 phosphorylation sites.**

AHL10 predicted structure indicates that in the tertiary structure of AHL10 the phosphorylation sites at S313 and S314 in the C-terminal loop region are not adjacent to the PPC/DUF296 domain involved in AHL multimerization nor the AT-hook domains involved in DNA binding.

AHL10 structure was predicted by I-Tasser and visualized by PyMOL. Color coding: Helices: Cyan, Loops: Wheat (pink), AT-hook1: Red surface, AT-hook2: Yellow surface PPC/DUF296 domain: magenta surface.

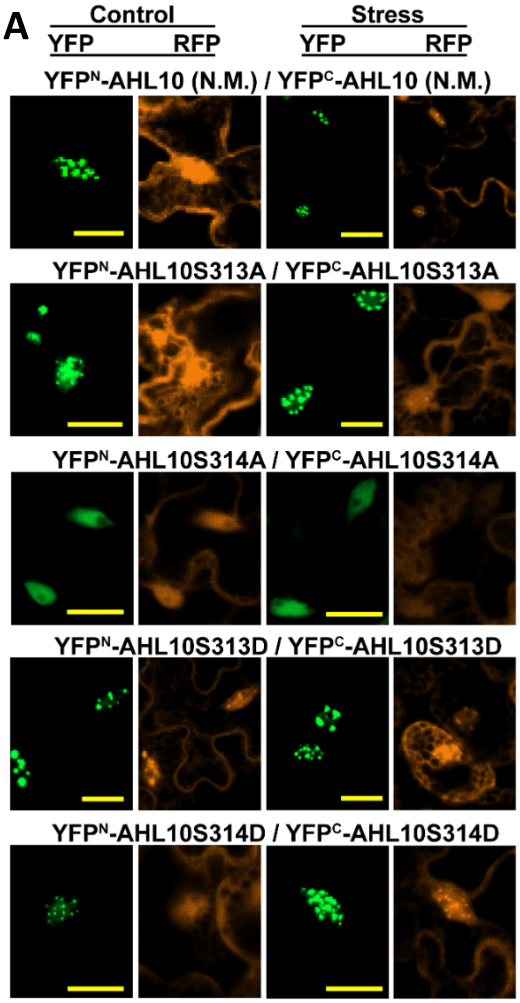


**SI Appendix Figure S13: Representative images of AHL10 self interaction in rBiFC assays.**

A. Representative images of AHL10 self-interactions (YFP) and constitutively expressed RFP used to normalize the interaction intensity. These images show representative examples of nuclei with AHL10 foci (except for S314A where nuclear foci were never observed). Scale bars indicate 20 μm.

B. Representative images of nuclei without foci for non-mutated AHL10 (N.M.) as well as the two phosphomimic AHL10 constructs. Scale bars indicate 20 μm.

Quantification of the proportion of nuclei with and without foci is shown in Fig. 4C.

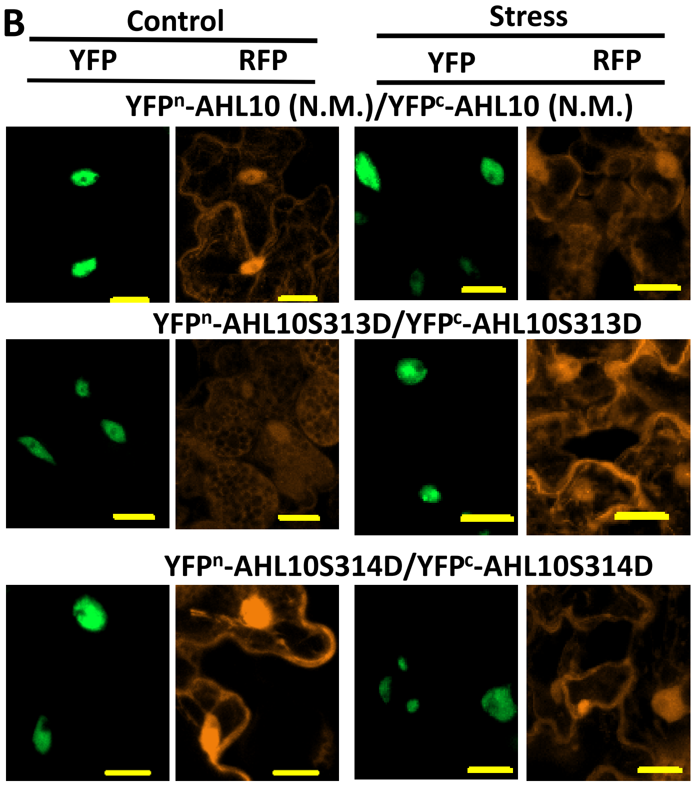

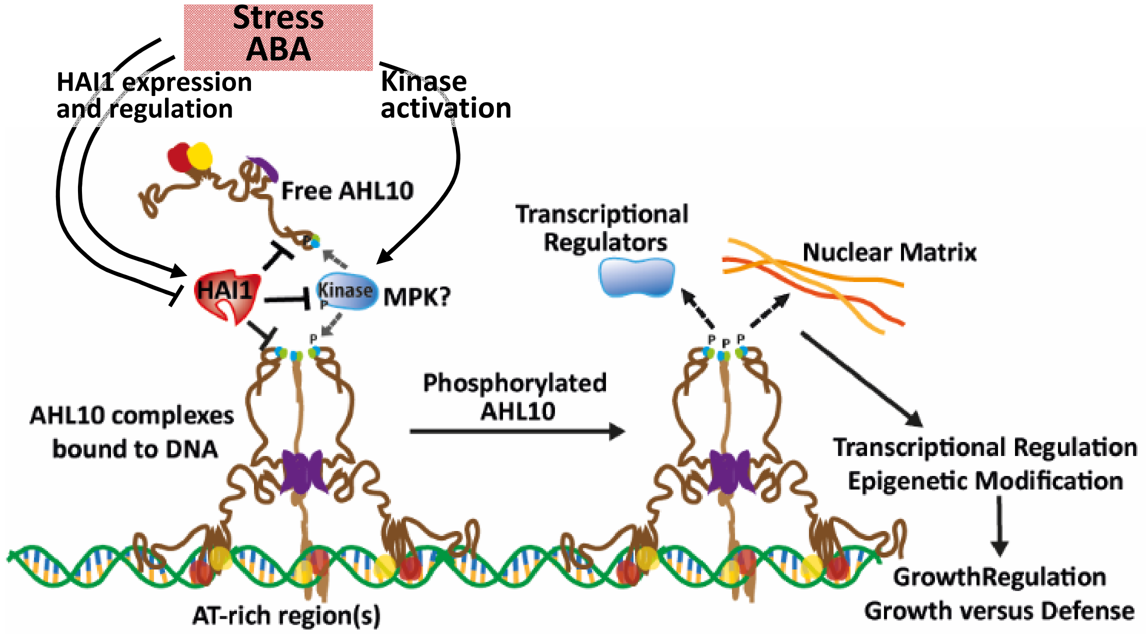


**SI Appendix Figure S14: Model of AHL10 function and its regulation by HAI1.** Previous analysis of AHLs indicates that they may bind DNA as trimers with each AHL in the complex binding either to adjacent DNA sites or to more distal sites to bring them together in the same regulatory complex. HAI1 dephosphorylates AHL10. This likely occurs on free AHL10 in the nucleoplasm (as the BiFC interactions of HAI1 and AHL10 do not show nuclear foci) but could also occur within DNA bound AHL complexes. HAI1 dephosphorylation is counteracted by phosphorylation by unknown kinase(s). HAI1 may also regulate AHL10 phosphorylation by dephosphorylating these kinase(s) which phosphorylate AHL10. The presence of an SP motif at the AHL10 S314 phosphorylation site, as well as previous finding that AHL13 is a putative MPK substrate, suggests phosphorylation by MPK(s). This is also consistent with known MPK functions in stress signaling and demonstration that HAI1 can dephosphorylate MPK3 and MPK6. Both HAI1 and kinase activity are regulated by environmental and hormone signals to influence AHL10 phosphorylation. AHL10 S314 phosphorylation does not affect the formation of AHL10 complexes (as indicated by rBiFC analysis) but does affect the localization of AHL10 complexes in nuclear foci. These foci may represent sites of nuclear matrix attachment as proposed for other AHLs; however, further investigation will be needed to establish the nature of AHL10 foci. The variable C-terminal region where the AHL10 S314 phosphorylation site is located has been proposed to be involved in AHL interaction with diverse types of transcriptional regulators. Thus, S314 phosphorylation may also affect AHL10 interaction with other transcriptional regulators. In either case, S314 phosphorylation is required for AHL10 function in transcriptional regulation, perhaps related to epigenetic mechanisms as previously found for AHL10. This leads to altered expression of a set of genes involved in AHL10-dependent growth regulation at low water potential and which may be more broadly involved in growth versus defense coordination.
