## Supplementary material for "Phosphoproteomics of Arabidopsis Highly ABA-Induced1 identifies AT-Hook Like10 phosphorylation required for stress growth regulation"

**Other supplementary materials for this manuscript include the following:**

Figs. S1 to S14, Datasets S1 to S10

### Quantitative phosphoproteomics using I-Traq labeling

Phosphoproteomic analysis of *hai1-2* was conducted in the same set of experiments as the analysis of *egr1-1egr2-1* (1) with the *hai1-2* and *egr1-egr2-1* samples collected simultaneously along with samples of wild type. Phosphoproteome data of all three genotypes has been deposited to the ProteomeXchange Consortium via the PRIDE partner repository with the data set identifier PXD004869. The wild type phosphoproteome data has also been submitted to PhosPhat (http://phosphat.uni-hohenheim.de/ ). Microarray analysis of *hai1-2* from the same set of experiments has been previously described (2) and is available on NCBI GEO under accession number GSE35258. In these experiments, seven day old seedlings were transferred to fresh half strength-MS agar plates (control) or PEG infused agar plates (stress, -1.2 MPa) for 96 h before sample collection. Samples were collected from three independent experiments (biological replicates) and the reproducibility of the stress treatment in each biological replicate was confirmed by measuring seedling proline content.

Sample preparation for phosphoproteomics generally followed previous protocols (3) with some modifications as described (1). In brief, after homogenation of samples in liquid nitrogen and trichoroacetic acid (TCA)/acetone solution (10% TCA in acetone containing 0.07% 2-mercaptoethanol), proteins were solubolized using dissolution buffer (9 M Urea, 25 mM Tris-HCl, pH 8.0) containing phosphatase inhibitors (Sigma cocktail 2 and 3). Six mg of total protein for each sample was reduced with 10 mM DTT, alkylated with 25 mM iodoacetamide. Protein digestion was performed by treating with Lys-C endopeptidase followed by trypsin. The tryptic peptides were desalted using a Sep-Pak C18 column and phosphopeptide enrichment performed using a Pierce $TiO_2$ phosphopeptide enrichment and clean-up kit. Phosphopeptide labeling was done using an AB Sciex iTRAQ 8-plex reagent kit following the manufacturer's instructions. iTRAQ reagents 113/117 were used for wild type control/stress and 115/119 for *hai1-2* control/stress. Samples labeled with individual iTRAQ tags were combined in one tube, desalted using a Pierce graphite spin column, desalted and fractionated by strong cation exchange (PolySULFOETHYL A column, 4.6 x 100 mm, 5 um, 200Å; PolyLC Inc. Columbia, MD) using an elution gradient from 0-50% buffer B (0.35 M KCl in buffer A; buffer B consisted of 7 mM KH2PO4, pH 2.65, 30% acetonitrile). Samples were eluted over 20 min and ten fractions of 0.5 ml each were collected. Each fraction was desalted using a graphite spin column and dried under vacuum.

The SCX fractions were analyzed on a nanoUPLC system (nanoAcquity, Waters, Milford, MA) coupled to an LTQ Orbitrap Velos hybrid mass spectrometer (Thermo Scientific,Bremen, Germany). A C18 capillary column (75 µm x 250 mm, 1.7 µm, BEH130, Waters, Milford, MA) was used to separate peptides with a 90-min linear gradient from 5% to 40% acetonitrile in 0.1% formic acid at a flow rate of 300 nl/min. The operation parameters for the LTQ Orbitrap Velos MS were as previously described (1). Peptide identification, phosphorylation, and quantification were performed using Proteome Discoverer software (v1.3, Thermo Fisher Scientific) with SEQUEST and Mascot (v2.3, Matrix Science) search engines. MS data were searched against TAIR10 protein sequence database with 27,416 sequence entries downloaded from the Arabidopsis Information Resource website (http://www.arabidopsis.org/). The parameters for database searches were set as follows: full trypsin digestion with 2 maximum missed cleavage sites, precursor mass tolerance = 10 ppm, fragment mass tolerance = 0.8 Da (CID) and 50 mmu (HCD), dynamic modifications: oxidation (M) and phosphorylation (S, T, and Y), static modifications: carbamidomethyl (C), iTRAQ 8-plex (N-terminus and K). The identified peptides were validated using Percolator algorithm which automatically conducted a decoy database search and rescored peptide spectrum matches (PSM) using *q*-values and posterior error probabilities. All the PSMs were filtered with a *q*-value threshold of 0.01 (1% false discovery rate). Phosphopeptides were further validated using phospho*RS* algorithm with thresholds of pRS score ≥ 50 and pRS site probability ≥ 75%. For iTRAQ quantification, the ratios of iTRAQ reporter ion intensities in MS/MS spectra (m/z:113 and 117) from raw datasets were used to calculate the fold changes in phosphorylation between control and treatment. Phosphopeptide abundances were statistically analyzed using a one sample T-test. To estimate the false discovery rates associated with our dataset, q-values were calculated from the characteristics of the p-value distribution. We estimated q-values using "*qvalue*" package in the R environment (4). The PhosPhat database http://phosphat.uni-hohenheim.de/ (5) was used to find additional experimentally identified AHL10 phosphorylation sites.

**Sequence Analysis and Structural Prediction**

Enrichment of specific motifs was analyzed using the Motif-X algorithm (6) ( http://motif-x.med.harvard.edu/ ). Sequences were manually centered at the phosphorylated residue and extended 10 amino acids on each side to generate pre-aligned 21 amino acid sequences (Dataset S4). The analysis parameters used were: minimum number of occurrences 5, significance 0.01, Arabidopsis proteome background. Alignment of Clade B AHL amino acid sequences was conducted using T-Coffee (http://www.tcoffee.org/ ) using the default parameters. Clade B AHL protein sequences were retrieved from Uniprot. BLOcks Substitution Matrix (BLOSUM62) was used to score sequence alignment of proteins and phylogenetic tree generated using average distance algorithm. Multiple sequence alignment of Clade B AHLs was done using Jalview (7). Structure prediction of AHL10 was performed using I-Tasser (8) and visualized in PyMol.

**Plant material, growth condition and physiological assays**

A T-DNA insertion line in AHL10, *ahl10-1* (SALK_073165C), was obtained from the Arabidopsis Biological Resource Center and the homozygous plants confirmed using primers designed using the Signal web resource; http://signal.salk.edu/. The *hai1-2* mutant (SALK_108282) has been previously described (2). To express *AHL10* under control of its native promoter, a genomic fragment containing the *AHL10* gene body and 2.1 kb of promoter sequence was amplified from Col-0, cloned into pENTR/D-TOPO (Invitrogen), moved to binary vector pGWB540 (9) and transformed into *Agrobacterium tumefaciens* strain GV3101 and then used for floral dip transformation of *ahl10-1*. For cDNA expression, cDNA was prepared from total RNA and the AHL10 cDNA amplified, cloned into pCR8/GW/TOPO (Invitrogen) followed by LR reaction into pEarleyGate104 (10) and transformation into *ahl10-1*. Substitution of Ser-313 (codon AGT) to Ala (codon GCT) or Asp (codon GAT) and Ser-314 (codon AGC) to Ala (codon GCC) or Asp (codon GAC) to generate phosphonull or phosphomimic of AHL10, respectively, was performed by PCR-driven overlap extension (11) using site-directed mutagenic primers and PCR product then transformed into pCR8/GW/TOPO. Plasmids isolated from spectinomycin-resistant colonies were sequenced to confirm the presence of the AHL10 mutation were then transferred by LR reaction to pEarleyGate104. The *hai1-2* mutant was introduced into *AHL10pro*:*AHL10-EYFP/ahl10-1* by crossing. For overexpression of HAI1, the HAI1 cDNA inserted into pEarleyGate 202 (2) was used to transform *AHL10pro*:*AHL10-EYFP/ahl10-1*. All constructs were verified by sequencing. Primers used for genotyping, cloning and site-directed mutagenesis are given in Dataset S10.

Seedling growth and low water potential treatment on PEG-infused agar plates was performed as described previously (1, 12). Growth chamber conditions were 22°C with continuous light at 110-130 µmol photons $m^{-2}$ $s^{-1}$. For seedling growth assays, five-day-old seedlings were transferred to fresh half-strength MS plates (control, -0.25 MPa) or PEG-infused plates (-0.7 MPa or -1.2 MPa; note that plates were always prepared and infused with PEG for 15 h before use to avoid drying of the media and generate plates of consistent water potential). Root length and fresh weight were measured five days after transfer for control and ten days after transfer for stress treatment. To quantify seedling weight, groups of seedlings were weighed to obtain fresh weight (4-6 seedlings per data point for the unstressed control, 6-7 seedling per data point for the -0.7 MPa treatment and 8-9 seedlings per data point for the -1.2MPa treatment), dried at 65°C and reweighed to obtain dry weight. Wild type seedlings were included on every plate and the growth of each mutant or transgenic line was calculated relative to wild type grown on the same plate to minimize the effect of plate to plate variability in the data. For experiments involving transgenic lines, three independent lines were evaluated for each construct and each line was measured in at least three independent experiments and two or three plates per experiment. Note that in Figure 2, the data from three independent transgenic lines for each construct has been combined. For experiments involving protein or gene expression assays seven-day-old seedlings were transferred to control or stress treatments and samples collected 96 h after transfer.

Controlled soil drying experiments were conducted as previously reported (13). A standard potting mix was combined with 25% Turface (Turface MVP, Profile Products LLC, USA). Four genotypes were planted in sectors (two plants per sector) of 8 cm x 8 cm x 10 cm (LxWxH) plastic pots to ensure that the different genotypes were exposed to the same extent of soil drying. Plants were grown in a short day chamber (8 h light period, 23 C, light intensity of

100-120 μmol $m^{-2}$ $sec^{-1}$) and Hyponex nutrient solution (1 g $liter^{-1}$) supplied once per week. The position of individual pots within the chamber was rotated every two or three days. On day 19 after planting, pots were watered to saturation, allowed to drain and weighed. Water was withheld for 12 days (leading to 50-60 percent reduction in pot weight), each pot re-watered to 75 percent of the initial pot weight and then allowed to dry another 8-10 day until pot weight again reached 50-60 percent of the starting weight. At the end of the experiment, representative rosettes were photographed and the rest weighed (fresh weight) and then dehydrated in an oven to obtain the dry weight.

**Subcellular localization and analysis of protein interaction by ratiometric Bi-molecular Fluorescence Complementation (rBiFC).**

Subcellular localization of AHL10 was analyzed in root cells of $T_3$ homozygous lines. YFP was detected using excitation wavelength of 514 nm at 70% intensity and emission filter of 520-555 nm band pass on a Zeiss LSM 510 META microscope. Ratiometric BiFC (rBiFC) assays were performed using previously described vectors (14). Coding sequences of AHL10 (or AHL10 phosphonull and phosphomimic) and HAI1 (or HAI2, HAI3) were cloned into pDONR221 P1-P4 and pDONR221 P3-P2 (Invitrogen) respectively (primers used are listed in Dataset S10). These donor vectors were used for LR reaction with the destination vector pBiFCt-2in1-NN. The resulting plasmid was transformed into *Agrobacterium tumefaciens* strain GV3101 and transient expression was performed in 5-day-old seedlings of a line with Dexamethasone (DEX) inducible AvrPto expression (ABRC stock number CS67140) (15). DEX application and Agrobacterium infiltration of seedlings was performed as previously described (16). Seedlings were transferred to fresh control or PEG-infused agar plates (-1.2 MPa) 24 h after infiltration and rBiFC interaction was visualized 48-96 h after transfer. To visualize the rBiFC interaction, YFP was detected on a Zeiss LSM 510 META microscope at excitation wavelength 514 nm (70-80% intensity) and emission filter of 520-555 nm band pass with 450-550 μm pinhole while RFP was detected at excitation wavelength 543 nm (70-80% intensity) and emission filter 560-615 nm band pass with 450-550 μm pinhole. To quantify the rBiFC ratio, fluorescence intensity of YFP and the constitutively expressed RFP reporter was quantified for at least 20 individual cells from 4-5 seedlings combined from at least two independent experiments. For each quantification, a whole cell was selected as a region of interest (ROI) in Image J, the mean intensity of RFP and YFP over the ROI determined (using the analyze/measure function) and the YFP/RFP ratio calculated. Given the emission spectra of YFP and RFP and the filter set used to detect emission, a low level of bleed through from the YFP channel to RFP is unavoidable. In pixels where YFP intensity was very high, this bleed through could be seen in the RFP images. Because this occurred in only a small number of pixels while the rBiFC analysis used RFP intensity over the whole cell, the YFP/RFP ratio used for quantitation of relative interaction intensities was not substantially affected by leakage between the two channels. Also, inspection of pixel by pixel scatter plots of YFP versus RFP intensities showed that leakage from the RFP channel into the YFP channel was minimal and did not affect the visualization of YFP localization. For rBiFC experiments of AHL10 self-interaction, the number of nuclei having two or more visible YFP foci were counted and nuclei not meeting this criteria were considered as having diffuse localization. For the counts of nuclei with AHL10 foci, 95% confidence intervals and significant differences based on Fisher's exact test were determined using GraphPad Quick Calcs ( https://www.graphpad.com/quickcalcs/catMenu/ ).

Additional BiFC analyses were performed in similar manner using pSITE-nEYFP-C1 and pSITE-cEYFP-C1vectors as previously described (1).

**Recombinant Protein Expression and Purification**

An N-terminal truncation (amino acids 1-103) of HAI1 was used for protein expression. Note that this truncation was similar to that used previously for in vitro expression and activity assays of HAI1 (17) and was necessary to improve HAI1 solubility. $HAI1^{\Delta 1-103}$ was cloned into pET300NT destination vector (N-terminal HIS tag) and transformed into the *E.coli* strain BL21 (Rosetta DE3). $HAI1^{\Delta 1-103}$ protein expression was induced by addition of 0.75 mM IPTG when cultures reached an optical density of 0.4. Following overnight incubation at 16 ºC, cells were harvested and resuspended in lysis buffer (50 mM Tris HCl pH 7.5, 250 mM KCl, 0.1% Tween-20, 10% glycerol, 10 mM β-ME, 1× Roche EDTA-free protease inhibitor) and lysed using cell disruptor (Constant Systems). Lysate was filtered and applied to a 5 mL Mini Profinity IMAC cartridge using a BioLogic low pressure chromatography system (Bio-Rad). An imidazole gradient was used to elute the protein in elution buffer **(**50 mM Tris HCl pH 7.5, 20% glycerol, 25 mM $Mg(OAc)_2$, 2 mM $MnCl_2$, 10 mM β-ME, 1X Roche EDTA-free protease inhibitor). Note that no additional salt was added to the eluted protein as salt interferes with phosphatase activity. Phosphatase activity of the recombinant $HAI1^{\Delta 1-103}$ was tested using a phosphatase assay kit (Biosciences Cat. #786-453).

***In vitro* dephosphorylation of AHL10 and analysis of AHL10 phosphorylation using Phos-Tag polyacrylamide gels.**

To obtain highly phosphorylated AHL10 protein for *in vitro* dephosphorylation assay, seedlings of *AHL10pro*:*AHL10-EYFP/ahl10-1hai1-2*, were collected after 96 h stress treatment, ground in liquid nitrogen and further homogenized in cold extraction buffer (150 mM NaCl, 50 mM Tris HCl pH 7.5, 10% glycerol, 10 mM EDTA, 10 mM DTT, 1mM NaF, 1mM $Na_2MoO_4$, 1% IGEPAL, 2× Roche EDTA-free protease inhibitor, 2× Roche phosphatase inhibitor). The homogenate was centrifuged and protein content of the supernatant measured using a Pierce 660nm Protein Assay Reagent (Thermo Scientific). For immunoprecipitation of AHL10-YFP, 30 µl of equilibrated GFP-Trap magnetic resin (Chromotek) was added to extract containing 100 µg of total protein and rotated at 4ºC for 3 h. The samples were maintained at 4ºC and the GFP-Trap resin washed four times with washing buffer (100 mM NaCl, 10 mM Tris HCl pH 7.5, 0.5% IGEPAL, 1mM DTT, 1X Roche EDTA-free protease inhibitor, 10% glycerol) followed by an additional wash and resuspension in phosphatase treatment buffer (10 mM Tris HCl pH 7.5, 10% glycerol, 1X Roche EDTA-free protease inhibitor). The GFP-Trap resin and bound protein was then equally divided into separate tubes for mock (no phosphatase) treatment, treatment with 1 µl calf intestinal (CIP, NEB) in 1× cut smart buffer or, treatment with 2.5 or 5.0 µg of purified HAI1 in elution buffer. The tubes containing immunoprecipitate plus CIP or HAI1 (or mock incubation without phosphatase) were incubated in 37ºC for 30 min and the reaction stopped by adding SDS loading buffer and heat inactivation for 10 min at 70°C. For additional analysis of AHL10 phosphorylation, *AHL10pro*:*AHL10-EYFP/ahl10*, *35S:YFP-AHL10* N.M. (not mutated), *35S:YFP-AHL10* S313A and *35S:YFP-AHL10* S314A, were stress treated for 96 h and immunoprecipitated following the protocol described above.

Preparation and use of $Mn^{2+}$-Phos-Tag polyacrylamide gels and subsequent immunoblot was performed as described previously (18) but with some conditions optimized for AHL10.

Phos-Tag gel was prepared using 10 ml of 8% acrylamide and 50 μM Phos-tag for the resolving gel [2.67 mL of 30% acrylamide/bis 37.5:1 (Biorad), 2.5 mL of 1.5 M Tris/HCl pH 8.8, 100 μL of 5 mM Phos-tag$^{TM}$ AAL (Wako), 100 μL of 10 mM $MnCl_2$, 100 μL of 10% SDS, 10 μL TEMED, 25 μL of 10% APS, 4.5 ml of distilled water] and 4 mL of 4.5% acrylamide for the stacking gel (600 μL of 30% acrylamide/bis 37.5:1 (Biorad), 1 mL of 0.5 M Tris/HCl pH 6.8, 40 μL of 10% SDS, 4 μL TEMED, 20 μL of 10% APS and 2.34 mL of distilled water). The gel was run initially at 80 V for 30 min at RT then transferred to 30 V for 16 h at 4°C using 1× running buffer (50mM Tris base pH8.3, 0.1% SDS, 192 mM glycine). The resolving gel was then soaked in 2× 100 mL of transfer buffer (192 mM glycine, 25mM Tris-base pH 8.0, and 10% methanol containing 10 mM EDTA) for 30 min each time and then washed once for 20 min in transfer buffer without EDTA. The gel was then further incubated in 100 mL transfer buffer containing 0.2% (w/v) SDS (two 20-min incubations). Protein was blotted onto PVDF membrane by wet tank procedure at 30 V (at constant voltage) for 16 h at 4°C. Membranes were blocked using 5% nonfat milk and probed with 1:2000 dilution of Roche Anti-GFP (Sigma-Aldrich Cat#. 11814460001) and 1:10000 dilution of anti-mouse HRP secondary antibody. Normal SDS-PAGE analysis and blotting (with HSC70 as loading control where appropriate) were conducted as previously described (1).

For detection of AHL10 *in vivo* phosphorylation status, seedlings were ground in liquid nitrogen and homogenized in extraction buffer (50 mM Tris HCl pH 7.5, 10% glycerol, 1mM DTT, 1mM NaF, 1% IGEPAL, 1× Roche EDTA-free protease inhibitor, 1× Roche phosphatase inhibitor) and protein content assayed as described above. Twenty-five μg of total protein was heated (70°C, 10 min) in SDS loading buffer and analyzed by $Mn^{2+}$-Phos-Tag polyacrylamide gel electrophoresis and immunoblotting as described above but with the Phos-Tag gel run for longer time period (20 h).

Immunoblot band intensities were quantified using the Fiji ImageJ package (http://fiji.sc/Fiji). An area of interest around the major band(s) in each lane was designated using the rectangular selection tool. The ‘plot lane’ option in the gel analyzer menu was used to obtain intensity peaks for each selected band. Peak areas for integration were marked, and peak areas calculated using the wand tool.

**RNA sequencing**

For RNA sequencing, 7-day-old seedlings of Col-0 wild type and *ahl10-1* were transferred to either fresh control agar plates or to PEG-infused plates (-0.7 MPa) and samples collected 96 h after transfer. Total RNA was extracted using RNeasy Plant Mini Kit (Qiagen) according to the manufacturer’s protocol and RNA quality checked by 25S/18S ratio using an Agilent Bioanalyzer 2100 according to manufacturer’s procedures. Three independent biological experiments were performed and one sample collected from each experiment and used for library preparation (12 RNAseq libraries prepared). The reproducibility of the stress treatment in each biological replicate was confirmed by measuring seedling fresh weight and dry weight. Library preparation and RNA sequencing was carried out by the High Throughput Genomics Core facility of the Biodiversity Research Center, Academia Sinica following protocols using 3 ug of total RNA for oligo-dT purification of mRNA and library construction by Illumina TruSeq Stranded Total RNA Library Prep Kit. USER digest treatment was performed for adaptor ligation and the final library was amplified by 10 cycles of PCR using KAPA HiFi Mastermix.

The sequencing was performed on an Illumina HiSeq 2500 using paired end reads with an average insert size of 250 bp. Base calling and demultiplexing were performed using Illumina CASAVA-Bcl2fastq version 1.8.4. RPKM values were calculated using the RackJ software package (http://rackj.sourceforge.net/) with reads mapped to the TAIR10 genome using Bowtie2 and BLAT with default parameters. Expression comparisons were performed using DESeq2.

**Quantitative Reverse Transcriptase-PCR analysis of gene expression**

Total RNA was extracted from control or -0.7 MPa stress-treated seedlings using RNeasy Plant Mini Kit (Qiagen). Total RNA (typically 1 μg) was reverse transcribed using SuperScript III (Invitrogen). Real-time quantitative PCR assays were prepared using KAPA SYBR FAST qPCR kit (Kapa Biosystems) and run on an Applied Biosystems QuantStudio 12K QPCR instrument. Gene expression values were calculated using the comparative cycle threshold method following normalization based on *ELF1α* expression. Primers used are given in Dataset S10.

**References**


1. Bhaskara GB, Nguyen TT, Yang TH, & Verslues PE (2017) Comparative Analysis of Phosphoproteome Remodeling After Short Term Water Stress and ABA Treatments versus Longer Term Water Stress Acclimation. *Frontiers in Plant Science* 8: 523
2. Bhaskara GB, Nguyen TT, & Verslues PE (2012) Unique Drought Resistance Functions of the Highly ABA-Induced Clade A Protein Phosphatase 2Cs. *Plant Physiol* 160(1):379-395.
3. Wu JA, *et al.* (2010) Integrating titania enrichment, iTRAQ labeling, and Orbitrap CID-HCD for global identification and quantitative analysis of phosphopeptides. *Proteomics* 10(11):2224-2234.
4. Storey JD, Bass AJ, Dabney A, Robinson D (2015). *qvalue: Q-value estimation for false discovery rate control*. R package version 2.12.0, http://github.com/jdstorey/qvalue.
5. Durek P, *et al.* (2010) PhosPhAt: the Arabidopsis thaliana phosphorylation site database. An update. *Nucleic Acids Research* 38:D828-D834.
6. Schwartz D & Gygi SP (2005) An iterative statistical approach to the identification of protein phosphorylation motifs from large-scale data sets. *Nature Biotechnology* 23:1391.
7. Waterhouse AM, Procter JB, Martin DMA, Clamp M, & Barton GJ (2009) Jalview Version 2—a multiple sequence alignment editor and analysis workbench. *Bioinformatics* 25(9):1189-1191.
8. Yang JY, *et al.* (2015) The I-TASSER Suite: protein structure and function prediction. *Nature Methods* 12(1):7-8.
9. Nakagawa T, *et al.* (2007) Improved Gateway binary vectors: high-performance vectors for creation of fusion constructs in transgenic analysis of plants. *Bioscience, Biotechnology, and Biochemistry* 71(8):2095-2100.
10. Earley KW, *et al.* (2006) Gateway-compatible vectors for plant functional genomics and proteomics. *Plant J* 45(4):616-629.
11. Heckman KL & Pease LR (2007) Gene splicing and mutagenesis by PCR-driven overlap extension. *Nature Protocols* 2(4):924-932.
12. Verslues PE, Agarwal M, Katiyar-Agarwal S, Zhu J, & Zhu JK (2006) Methods and concepts in quantifying resistance to drought, salt and freezing, abiotic stresses that affect plant water status. *Plant J* 45(4):523-539.
13. Bhaskara GB, Yang T-H, & Verslues PE (2015) Dynamic proline metabolism: importance and regulation in water limited environments. *Frontiers in Plant Science* 6: 484
14. Grefen C & Blatt MR (2012) A 2in1 cloning system enables ratiometric bimolecular fluorescence complementation (rBiFC). *Biotechniques* 53(5):311-314.
15. Tsuda K, *et al.* (2012) An efficient Agrobacterium-mediated transient transformation of Arabidopsis. *Plant J* 69(4):713-719.
16. Kumar MN, Hsieh YF, & Verslues PE (2015) At14a-Like1 participates in membrane-associated mechanisms promoting growth during drought in Arabidopsis thaliana. *Proc Natl Acad Sci USA* 112(33):10545-10550.
17. Antoni R, *et al.* (2012) Selective inhibition of clade A phosphatases type 2C by PYR/PYL/RCAR abscisic acid receptors. *Plant Physiol* 158(2): 970-980.
18. Kinoshita E, Kinoshita-Kikuta E, Takiyama K, & Koike T (2006) Phosphate-binding tag, a new tool to visualize phosphorylated proteins. *Molecular & Cellular Proteomics* 5(4):749-757.
